# Glycogen deficiency impairs diurnal energy metabolism and cell division in *Synechocystis*

**DOI:** 10.64898/2026.05.22.726976

**Authors:** Jan M. Hofer, Tim Schulze, Lennart Witting, Bianca Laker, Stephan Krueger, Philipp Westhoff, Dietrich Kohlheyer, Andreas P.M. Weber, Marion Eisenhut

**Author notes:** Corresponding author: Marion Eisenhut. Authors contributed equally (Jan M. Hofer, Tim Schulze). The author responsible for distribution of materials integral to the findings presented in this article in accordance with the policy described in the Instructions for Authors (https://academic.oup.com/plphys/pages/General-Instructions) is Marion Eisenhut.

## Abstract

Diurnal changes in light availability are a defining feature of life on Earth. Photoautotrophic organisms therefore store reduced carbon during the day to sustain energy metabolism at night. In cyanobacteria, glycogen is the primary carbon storage compound and supports both energy homeostasis and stress responses. Although glycogen-deficient *Synechocystis* strains have been studied previously, how these mutants cope with the loss of the major daytime carbon sink and can sustain themselves during the night remains unclear. Using single-cell microfluidics, transcriptomics, and metabolomics, we show that Δ*glgC* mutants exhibit pronounced light sensitivity. At sub-lethal light intensities, daytime transcriptional responses are dominated by downregulation of photosynthesis-related genes, likely preventing NADPH overaccumulation in the absence of a carbon sink. During the night, mutants display severe energy limitation, characterized by reduced ATP levels, altered redox balance, and depletion of central carbon intermediates. In contrast, fumarate and malate accumulate, indicating enhanced respiratory flux through succinate dehydrogenase. These metabolic constraints lead to extended lag phases and delayed cell divisions after the onset of light, demonstrating that glycogen-deficient cells fail to efficiently reinitiate growth after dawn.

Overall, our results as a snapshot of the initial response to diurnal regimes highlight glycogen as a central integrator of diurnal physiology in *Synechocystis*, coordinating energy metabolism, redox balance, and cell division, with implications for metabolic robustness and the evolutionary constraints shaping (endo)symbiosis.

**Short summary:** Glycogen deficiency disrupts day–night energy and redox homeostasis in *Synechocystis*, revealing constraints on growth, division, and symbiotic potential.

## INTRODUCTION

Cyanobacteria are photoautotrophic organisms that depend on solar energy for growth and survival. To cope with periods of darkness, they have evolved distinct strategies. While short-term light fluctuations during the day can be compensated by modulating energy fluxes around the photosystems (Mullineaux, 2014), the long-term absence of light during night requires a more extensive adaptation: the synthesis of polysaccharide storage compounds during the day that serve as carbon and energy reserves at night. Because diurnal light–dark cycles are ubiquitous for phototrophic organisms, such storage compounds are widespread across photosynthetic lineages. In the Archaeplastida, which acquired their plastids from a cyanobacterial ancestor via endosymbiosis (Rodríguez-Ezpeleta et al., 2005), two major storage polysaccharides predominate: glaucophytes and rhodophytes store floridean starch in the cytosol, whereas Viridiplantae store starch in plastids (Ball et al., 2011). In cyanobacteria, glycogen - a branched glucose polymer also found in many heterotrophic bacteria - is the dominant storage compound, although granulose and starch-like polymers have also been reported in unicellular diazotrophic strains to provide energy for nitrogen fixation, a process only occurring at night due to incompatibility with oxygenic photosynthesis (Nakamura et al., 2005; Deschamps et al., 2008; Beck et al., 2012; Suzuki et al., 2013; Colpaert et al., 2020). Whereas heterotrophic bacteria typically synthesize glycogen when organic carbon is abundant, but another nutrient is limiting (Preiss, 1984), glycogen plays a more central role in cyanobacterial metabolism due to the pronounced energetic constraints imposed by diurnal cycles (Angermayr et al., 2016; Cano et al., 2018; Shinde et al., 2020). During the light phase, a substantial fraction of fixed carbon is channeled into glycogen synthesis to ensure the availability of respiratory substrates during the night (Reimers et al., 2017; Lucius and Hagemann, 2024). Across organisms and storage compounds, these processes are often tightly regulated in response to light availability and circadian control (Lee et al., 2024).

In the cyanobacterial model organism *Synechocystis* sp. PCC 6803 (hereafter *Synechocystis*), four key enzymes catalyze glycogen synthesis, forming a pathway analogous to that of *Escherichia coli* (Preiss, 1984). Glucose 1-phosphate adenylyltransferase, encoded by *glgC* (*slr*1176), also termed ADP-glucose pyrophosphorylase (AGP), catalyzes the first committed step by activating glucose 1-phosphate through ATP-dependent adenylyl transfer. Subsequently, the glycogen synthases GlgA1 (Sll0945) and GlgA2 (Sll1393) elongate linear α-1,4-linked glucan chains (maltodextrins) using ADP-glucose as a substrate, releasing ADP. The branching enzyme GlgB (Sll0158) introduces α-1,6 linkages, thereby converting maltodextrins into branched glycogen. During the night, glycogen degradation is initiated by the glycogen phosphorylases GlgP1 (Sll1356) and GlgP2 (Slr1367), which cleave α-1,4 glycosidic bonds at the non-reducing ends, releasing glucose 1-phosphate. Because GlgP cannot hydrolyze α-1,6 linkages, the debranching enzyme GlgX (GlgX1, Slr1857, and GlgX2, Slr0237) is required to hydrolyze branch points, releasing maltotetraose. These oligosaccharides are subsequently processed by an α-1,4-glucanotransferase, which elongates acceptor chains while releasing free glucose. Finally, maltodextrin phosphorylase converts malto-oligosaccharides into glucose 1-phosphate, generating progressively shorter chains (Ball and Morell, 2003; Ball et al., 2011). Glucose 1-phosphate is then converted to glucose 6-phosphate by phosphoglucomutase (PGM, Sll0726), reconnecting glycogen catabolism to central carbon metabolism (Ortega-Martínez et al., 2023), from where carbon is primarily metabolized via the oxidative pentose phosphate pathway (OPPP) (Shinde et al., 2020). Glycogen-deficient *Synechocystis* mutants, including Δ*glgC* and Δ*glgA1*/Δ*glgA2* strains, have been studied extensively. These mutants display impaired growth under diurnal light regimes and altered responses to nitrogen starvation (Carrieri et al., 2012; Gründel et al., 2012; Hickman et al., 2013; Carrieri et al., 2017; Cano et al., 2018; Díaz-Troya et al., 2020; Ortega-Martínez et al., 2023), phenotypes that have also been observed in glycogen-deficient *Synechococcus elongatus* PCC 7942 strains (Suzuki et al., 2010; Hickman et al., 2013; Benson et al., 2016). In addition, mixotrophic growth (Carrieri et al., 2012; Gründel et al., 2012; Hickman et al., 2013; Carrieri et al., 2017; Díaz-Troya et al., 2020; Ortega-Martínez et al., 2024), as well as responses to oxidative and salt stress (Suzuki et al., 2010), are affected in these mutants. Although growth defects of glycogen-deficient strains under diurnal conditions have been consistently reported (Gründel et al., 2012; Cano et al., 2018; Shinde et al., 2020), the underlying mechanistic basis of these phenotypes remains insufficiently understood.

Cyanobacteria impaired in glycogen synthesis have also attracted considerable interest as potential production strains for biotechnological applications, because the absence of a major carbon sink can redirect carbon fluxes towards the synthesis of valuable products (Ducat et al., 2012; Li et al., 2014; Benson et al., 2016; Domínguez-Lobo et al., 2024; Kato et al., 2024).In *Synechococcus elongatus* PCC 7002, it has been suggested that up to 39% of photosynthetic capacity can be redirected towards the production of biotechnologically relevant compounds without compromising photoautotrophic growth (Xu et al., 2013).

The loss of glycogen synthesis also appears to represent a key evolutionary step during the transition from an endosymbiotic cyanobacterium to a plastid in the last common ancestor of the Archaeplastida (Rodríguez-Ezpeleta et al., 2005; Deschamps et al., 2008). During plastid evolution, the loss of autonomous carbon storage in the endosymbiont likely enforced metabolic interdependence with the host, a prerequisite for stable endosymbiosis and subsequent organellogenesis (Facchinelli et al., 2013; Karkar et al., 2015). This is consistent with the shift in carbon storage strategies within the Archaeplastida, where cyanobacterial glycogen storage was likely first relocated to the host cytosol before being replaced by starch in the Chlorophyta (Deschamps et al., 2008; Ball et al., 2015). The absence of glycogen in the more recently evolved chromatophores of *Paulinella* spp. (Nowack et al., 2008) further supports loss of glycogen storage and metabolic connectivity as a critical requirement for endosymbiont integration (Colleoni et al., 2010; Facchinelli et al., 2013; Karkar et al., 2015). Similarly, many intracellular pathogenic and symbiotic bacteria lack glycogen as a carbon storage compound (Henrissat et al., 2002).

In this study, we aimed to obtain a mechanistic understanding of how *Synechocystis* copes with glycogen deficiency under diurnal conditions. Using a recently developed microfluidic platform for the cultivation of phototrophic microorganisms (Witting et al., 2025), we identified pronounced phenotypic effects of glycogen deficiency on cellular physiology and morphology. To further dissect the underlying mechanisms, we performed comparative transcriptomic and metabolic analyses. Our results reveal that Δ*glgC* mutants experience acceptor limitation during the light phase and an ATP shortage during darkness. The mutants partially compensate for the loss of a carbon sink by restructuring cellular metabolism and machinery, leading to a reduction in photosynthetic capacity at the transcriptomic level. During the dark phase, alternative pathways are activated to generate NADPH and ATP at the expense of metabolites required to efficiently reinitiate carbon fixation during the dark-to-light transition.

## RESULTS

To understand the effects of the loss of the carbon storage compound glycogen in *Synechocystis*, we generated knock-out mutants in the *glgC* gene (*slr*1176), which encodes ADP-glucose pyrophosphorylase (AGP), the enzyme catalyzing the first step of glycogen biosynthesis. After multiple rounds of re-streaking on antibiotic-containing BG11 plates, we obtained two independent, fully segregated mutant clones, Δ*glgC*-1 and Δ*glgC*-6 (Figure 1A). Quantification of glycogen content confirmed the complete absence of glycogen synthesis in both mutants (Figure 1B).

**Figure 1.**
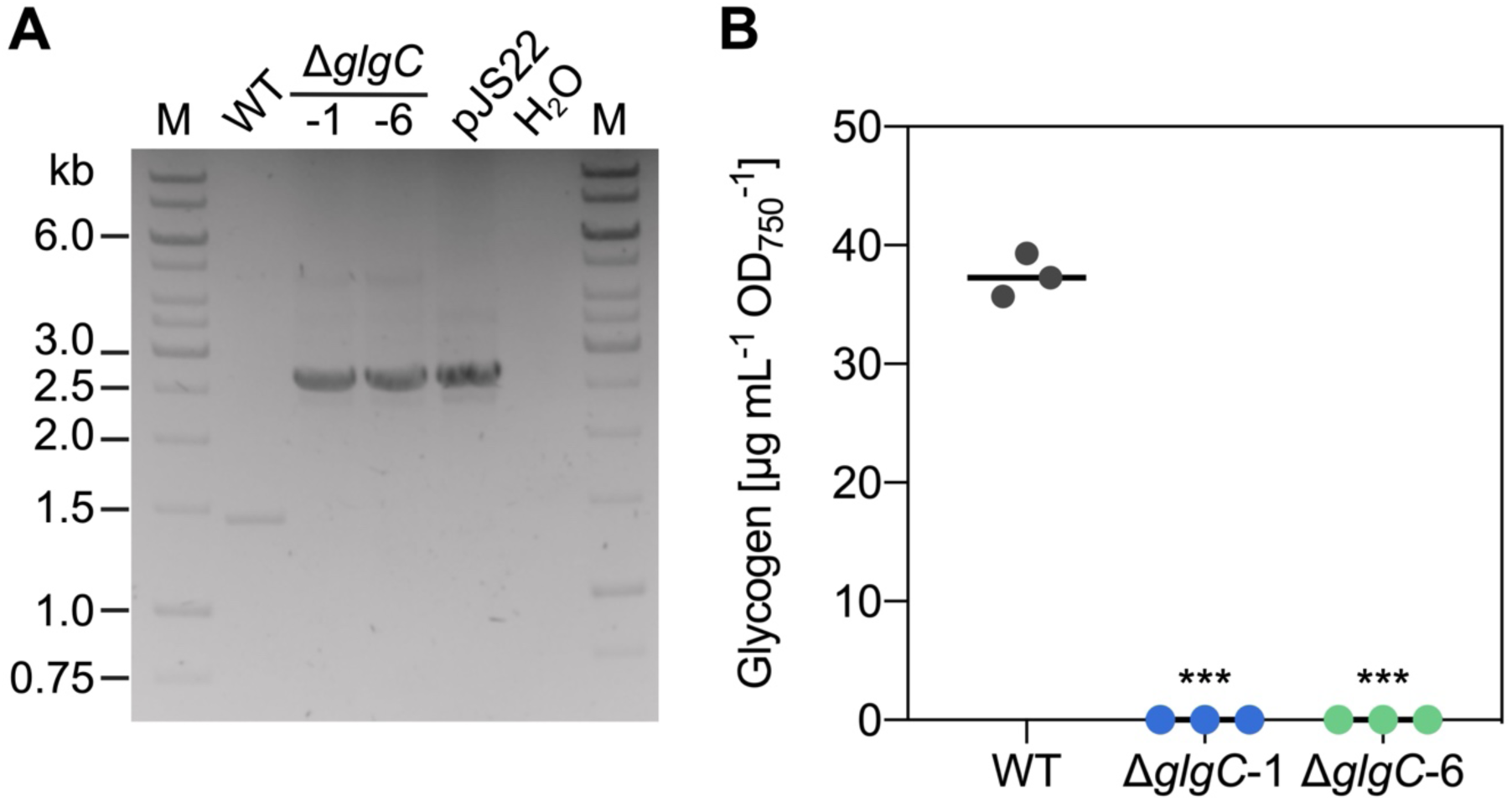
Generation of *Synechocystis* Δ*glgC* mutants. (A) Genotyping of Δ*glgC*-1 and Δ*glgC*-6 using genomic DNA and primers JS53 and JS54. Expected fragment sizes were 1,450 bp for the wild type (WT) and 2,700 bp for Δ*glgC*. Plasmid pJS22, containing the homologous recombination template, served as positive control; water (H_2_O) was used as negative control. M: molecular marker. (B) Intracellular glycogen content in WT, Δ*glgC*-1, and Δ*glgC*-6 cells grown in continuous light (100 µmol photons m^−2^ s^−1^) at 30 °C. Data represent mean and individual values of three biological replicates. Asterisks denote statistical significance between genotypes as calculated with the Student’s t-test (*\*\*\**: *p ≤* 0.001).

### Glycogen-deficient mutants are light-sensitive and show delayed growth resumption after darkness

To investigate the effect of glycogen loss on growth performance, we employed a recently established microfluidic cultivation for cyanobacteria, including *Synechocystis* (Witting et al., 2025). This approach enables cultivation of cells in single-cell layers within microfluidic chambers, allowing time- and spatially-resolved observation of individual cells. We first analyzed light-intensity-dependent growth (Figure 2A, B). Wild-type (WT) cells exhibited growth over the whole range of tested light intensities, with optimal growth at 80 µmol photons m^−2^ s^−1^. In contrast, both Δ*glgC* mutant lines grew comparably to the WT up to 35 µmol photons m^−2^ s^−1^ (Figure 2A) but failed to grow at higher light intensities (Figure 2B), indicating pronounced light sensitivity. To demonstrate the difference between growing and non-growing colonies, videos of Δ*glgC*-1 growing in the microfluidic device at light-intensities of 11 (Supplementary Video 1) and 56 µmol photons m⁻² s⁻¹ (Supplementary Video 2) are included in the supplementary material. Consequently, all subsequent microfluidic experiments were conducted at 30 µmol photons m^−2^ s^−1^.

**Figure 2.**
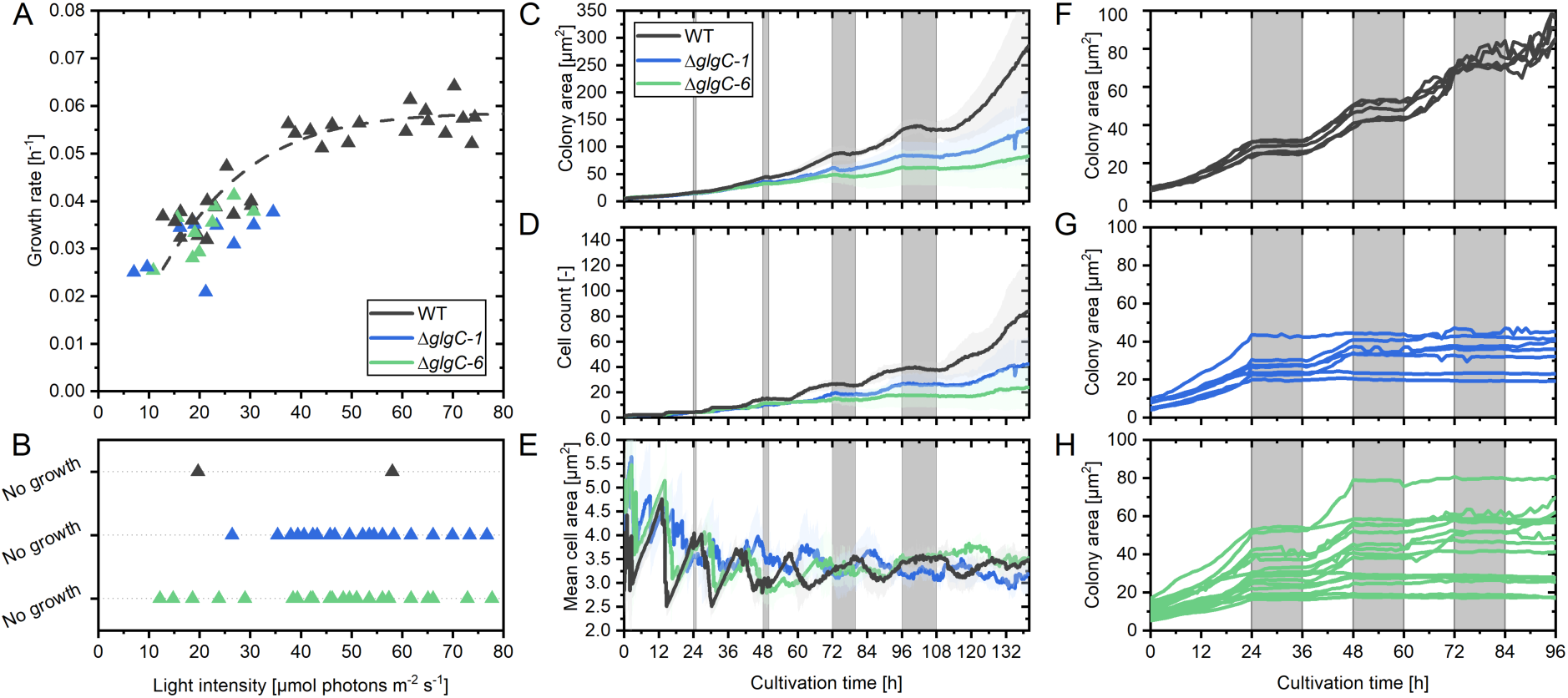
Phenotypic characterization of *Synechocystis* WT and Δ*glgC* mutants in a microfluidic chemostat. (A-B) Growth rates of WT and Δ*glgC* strains across the applied light intensity gradient. Each point represents one growth chamber. All strains were tested in two independent experiments. The same light-intensity gradient was applied to all strains. Observed light-dependent growth rates are given in (A). A hyperbolic tangent fit was applied to the WT data. (B) shows observed cells that did not display growth at given light intensities. (C–E) Time-resolved measurements of cumulative colony area (C), cell count (D), and mean single-cell area (E) under sequential light-dark cycles. Light periods were illuminated at 30 µmol photons m^−2^ s^−1^. Night phases without illumination are indicated in grey. Data represent mean ± SD from three independent experiments. (F–H) Growth under 12-h light/dark diurnal conditions over 96 h. Light periods were illuminated at 30 µmol photons m⁻² s⁻¹. Night phases without illumination are indicated in gray. Each line represents growth within a distinct growth chamber. The panels include five, seven, and 15 replicates for the WT (F), Δ*glgC-1* (G), and Δ*glgC-6* (H) mutants, respectively.

Next, we examined growth behavior under diurnal light–dark cycles. We hypothesized that growth impairment caused by glycogen deficiency would increase with longer dark periods and would result in full arrest after few consecutive 12-h light/dark cycles. Figures 2C–E show an experiment in which cells were cultivated for 140 h under light–dark cycles with increasing night lengths (1, 2, 6, and 12 h). Analysis of colony area and cell number over time revealed that WT cells performed best under all conditions. Although no cell division occurred during darkness in any strain, WT cells increased in size at the beginning of the dark phase (Figure 2C, Close-up images for 1-h and 2-h nights in Supplementary Figure S1), we refer to as nocturnal swelling. In contrast, Δ*glgC* mutants did not show nocturnal swelling and consistently required more time to resume growth after the dark phase, an effect particularly evident in colony area measurements (Figure 2C, Supplementary Figure S1). Following the 12-h night phase, WT cells resumed growth immediately, whereas Δ*glgC* mutants exhibited a delayed transition to exponential growth. During this final phase, growth rates of the mutants were lower than those of the WT. The growth rate of Δ*glgC*-1 was µ = 0.018 ± 0.006 h^−1^, for Δ*glgC*-6 µ = 0.013 ± 0.007 h^−1^, and µ = 0.029 ± 0.010 h^−1^ for the WT. Time-lapses of representative growth phenotypes in microfluidic chambers for each strain can be found in Supplementary Videos 3-5. We note that the successive light and dark periods are part of a continuous experimental sequence and therefore not independent conditions, as the physiological state of the cells is influenced by preceding phases. In an independent experiment we tested for how long the mutants could survive consecutive 12-h light/dark cycling (Figure 2F-H). In all WT chambers (Figure 2F), growth continued unaffected after the third consecutive 12-h night. For the Δ*glgC*-1 (Figure 2G) and Δ*glgC*-6 (Figure 2H) mutants we observed different growth behavior for individual colonies. Some colonies stopped growing immediately after the first 12-h dark period, some arrested after the second night, and only very few (one Δ*glgC*-1 colony and two Δ*glgC*-6 colonies) resumed growing after three nights. Importantly, the observation in the microfluidics platform demonstrated that the mutant cells were actually not lysing but arrested in growth.

Based on these phenotypes, we aimed at taking a snapshot of the initial response of the glycogen-deficient *Synechocystis* mutants to diurnal regimes, before effects of glycogen loss lead to full growth arrest. We hypothesized that glycogen loss in Δ*glgC* mutants affects photosynthesis, cellular energy status, and cell division control. To test these hypotheses and identify the underlying mechanisms, we next analyzed transcriptomic and metabolomic responses in batch cultures under controlled diurnal conditions. This approach is complementary to the single-cell phenotypes observed in microfluidics and allows to harvest sufficient cell material for these analyses. Using the conversion factor of 4.6 between light intensity in the microfluidic setup and the apparent light intensity in our batch cultivation system, Multi-Cultivator (Witting et al., 2025), we concluded that cultivation in the Multi-Cultivator device should be conducted below a light intensity of 138 µmol photons m⁻² s⁻¹ to be non-lethal for Δ*glgC* mutants. Thus, we performed all batch cultivation using a Multi-Cultivator device with 100 µmol photons m^−2^ s^−1^ as light intensity.

### Glycogen deficiency has a stronger impact in the dark than in the light

For transcriptomic and metabolomic profiling, the experimental design (Figure 3A) included a 36-h acclimation period with one 12-h light/dark cycle to entrain cells previously grown under continuous illumination. Samples were collected during a subsequent diurnal cycle. End-of-day (EoD) samples were taken after 11 h of illumination to capture glycogen accumulation in the WT and the consequences of abolished glycogen synthesis in the mutants during the light phase. End-of-night (EoN) samples were taken after 11 h of darkness to capture metabolic and transcriptional states associated with glycogen deficiency during the dark phase, as observed in microfluidic experiments. The experiment was conducted in two independent runs, each comprising two separately inoculated cultures per strain derived from a common pre-culture and cultivated independently under identical conditions, yielding a total of four biological replicates per condition.

**Figure 3.**
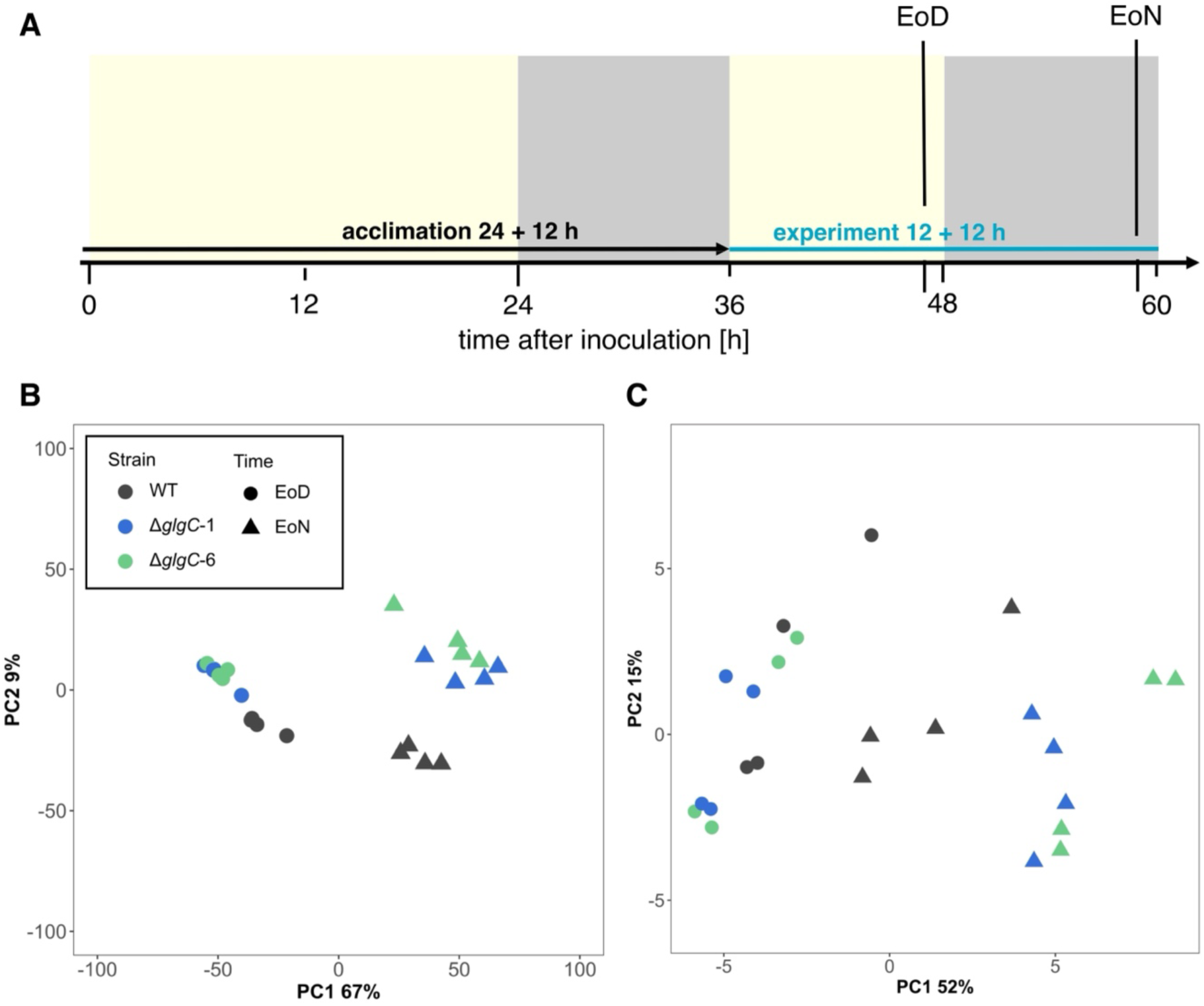
Overview of transcriptomic and metabolomic responses in WT and Δ*glgC* mutants. (A) Schematic of cultivation and sampling for end-of-day (EoD) and end-of-night (EoN) time points. Light periods are shaded yellow, dark periods in grey. Cultivation was performed in a Multicultivator at 30 °C, 100 µmol photons m^−2^ s^−1^, with ambient air bubbling. (B) Principal component analysis (PCA) of transcript abundances in WT, Δ*glgC*-1, and Δ*glgC*-6. Transcript per million (TPM) values were used for the analysis. (C) PCA of metabolite abundances in WT and Δ*glgC* mutants.

Principal component analysis (PCA) of transcriptomes (Figure 3B), based on transcripts per million (TPM) values, revealed four distinct clusters separating samples by sampling time and genotype. Combined, PC1 and PC2 explained 76% of the total variance, with PC1 alone accounting for 67%. Separation along PC1 clearly distinguished EoD from EoN samples, whereas PC2 (9% variance) separated samples by genotype. This indicates that diurnal conditions exert a stronger influence on transcript abundance than glycogen deficiency *per se*.

A comparable PCA of relative metabolite abundances (Figure 3C) explained 67% of the total variance, with PC1 and PC2 contributing 52% and 15%, respectively. Separation between EoD and EoN samples occurred along both components. Genotype-dependent divergence was apparent at EoN but not at EoD, consistent with the transcriptomic data.

### *Synechocystis* mounts a diurnal transcriptional response

Comparison of WT transcriptomes between EoD and EoN revealed extensive diurnal reprogramming, consistent with the PCA results (Figure 3B). Fifty-four percent of all genes were differentially expressed, with 1,171 genes (32%) significantly downregulated and 803 genes (22%) significantly upregulated at EoN (*q* ≤ 0.01; Figure 4A). GO term enrichment analysis of differentially expressed genes (DEGs) (Figure 4B) showed that transcripts related to photosynthesis, including light reactions, pigment and porphyrin biosynthesis, and carbon fixation (e.g., glyceraldehyde 3-phosphate metabolism), were predominantly downregulated at EoN. Additional metabolic processes, such as fatty acid, nucleoside phosphate, ribonucleoside triphosphate biosynthesis, and purine ribonucleotide metabolism were also reduced. Enriched GO terms among downregulated genes further included “nitrogen compound transport” and “proton transmembrane transport”.

**Figure 4.**
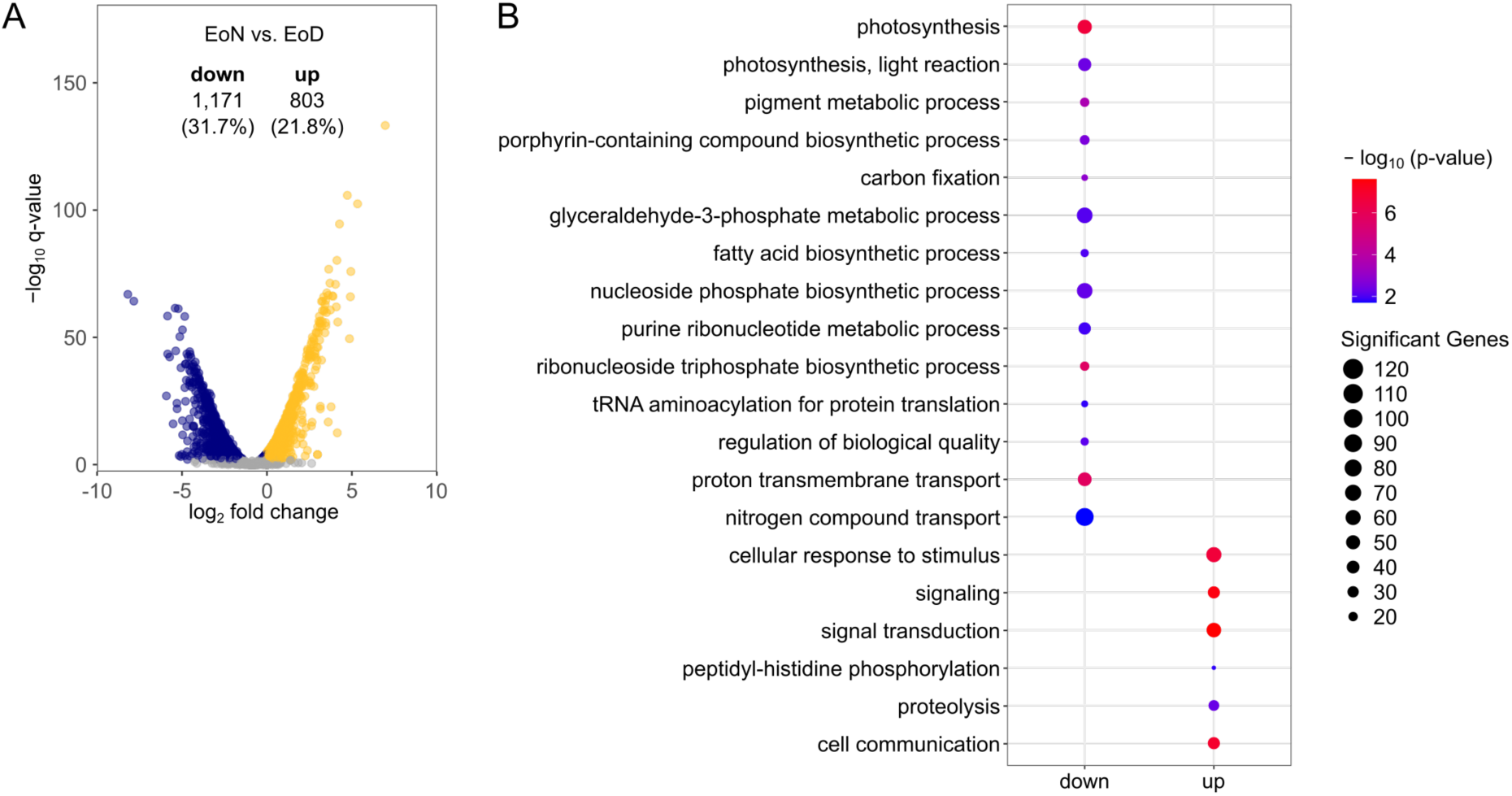
Global transcriptional changes in WT at EoN versus EoD. (A) Volcano plot of differentially expressed genes (DEGs, *q* ≤ 0.01) comparing EoN to EoD. Numbers indicate up-and downregulated genes and their percentage of total coding genes (3,691). (B) GO term enrichment of DEGs (*q* ≤ 0.01) in WT cells. Top 20 biological process GO terms with *p* ≤ 0.05 are shown.

In contrast, genes involved in “cellular responses to stimuli”, including “signal transduction”, were generally upregulated at EoN. The GO term “proteolysis” was also enriched, indicating increased protein turnover during the dark phase.

This diurnal transcriptional response (Supplementary Dataset S1) was generally also mounted by both Δ*glgC* mutant lines. Quantity (Supplementary Figure S2A, B) and quality (Supplementary Figure S2C) of the changes were comparable to the WT. GO term enrichment showed congruent results for downregulated transcripts in the categories “photosynthesis”, including “light reactions”, pigment and porphyrin biosynthesis, and “carbon fixation”. Upregulated transcripts were likewise enriched in the categories “cellular responses to stimuli”, including “signal transduction” (Supplementary Figure S2C).

### Glycogen deficiency elicits distinct transcriptomic responses at end of day and end of night

To assess genotype-specific effects, we analyzed DEGs and enriched GO terms separately for EoD and EoN. At EoD, Δ*glgC*-1 exhibited 174 downregulated (4.7%) and 218 upregulated genes (5.9%) (Figure 5A). In Δ*glgC*-6, 97 genes (2.6%) were downregulated and 145 genes (3.9%) upregulated (Figure 5B). GO term enrichment revealed shared downregulation of “photosynthesis”, “light reaction”, and “iron–sulphur cluster assembly”, whereas upregulated genes were enriched for “lipopolysaccharide biosynthetic process”, “inorganic anion transmembrane transport”, and “monoatomic anion transmembrane transport” (Figure 5C). At EoN, genotype-dependent differences were more pronounced. Compared to WT, Δ*glgC*-1 showed 281 downregulated (7.6%) and 250 upregulated genes (6.8%) (Figure 5D), while Δ*glgC*-6 displayed 316 downregulated (8.6%) and 305 upregulated genes (8.3%) (Figure 5E). Downregulated genes were enriched for “photosynthesis”, “light reaction”, “proton transmembrane transport”, and “protein catabolic process” (Figure 5F). Upregulated genes showed enrichment for “nicotinamide nucleotide biosynthetic process”, “amide biosynthetic process”, and “cell wall organization or biogenesis”.

**Figure 5.**
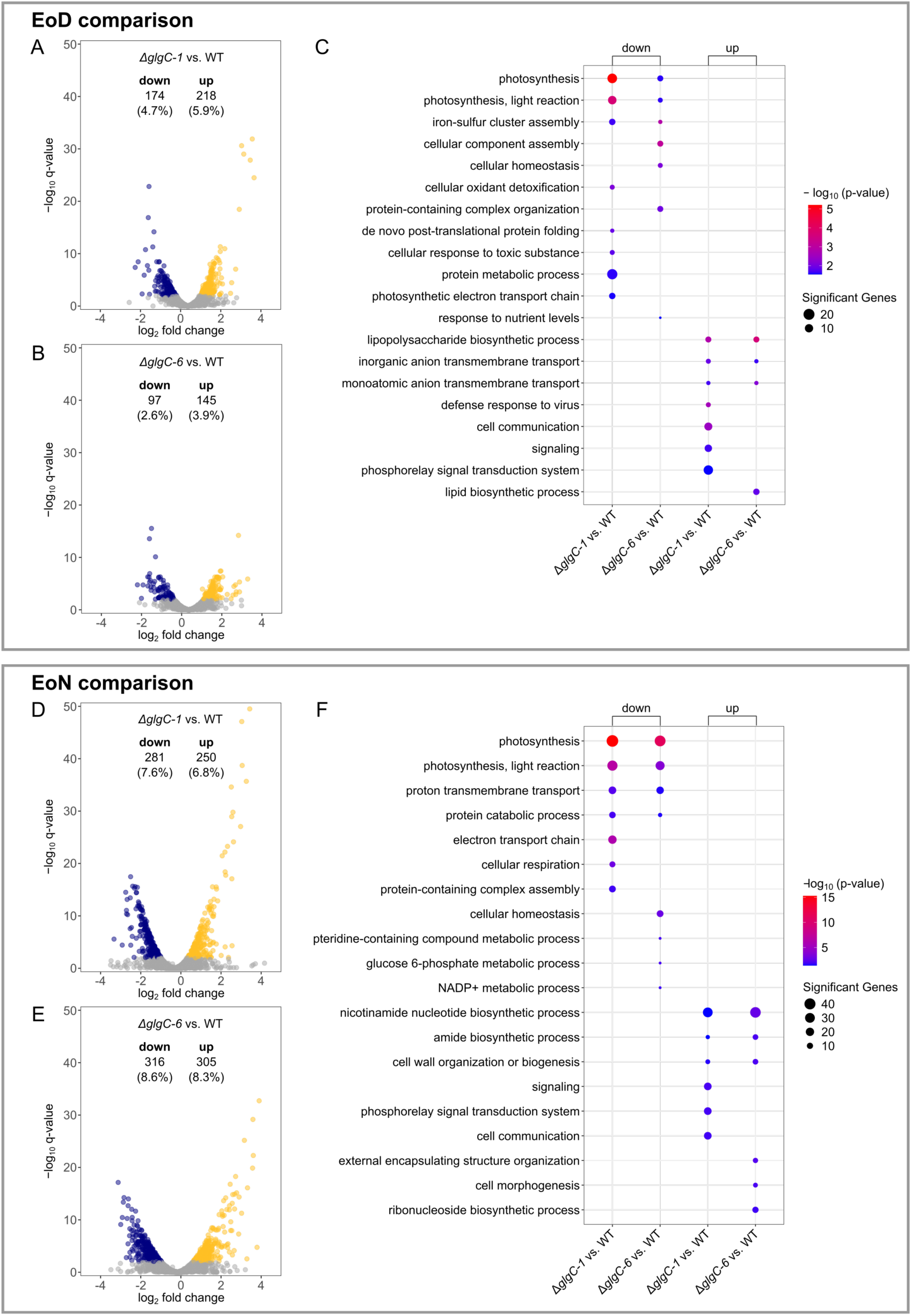
Transcriptional responses of Δ*glgC* mutants compared to WT. (A) Volcano plot of DEGs (*q* ≤ 0.01) in Δ*glgC*-1 versus WT at EoD. For all Volcano plots, numbers of significantly up- and downregulated genes and their percentage of total coding genes (3,691) are given. (B) Volcano plot of DEGs (*q* ≤ 0.01) in Δ*glgC*-6 versus WT at EoD. (C) GO term enrichment of DEGs (*q* ≤ 0.01) for both Δ*glgC* lines at EoD. Shown are the TOP20 of enriched GO terms (biological processes) with a *p*-value ≤ 0.05. (D–E) Volcano plots of DEGs (*q* ≤ 0.01) in Δ*glgC*-1 (D) and Δ*glgC*-6 (E) versus WT at EoN. (F) GO term enrichment of DEGs (*q* ≤ 0.01) at EoN for both Δ*glgC* mutants. Shown are the TOP20 of enriched GO terms (biological processes) with a *p*-value ≤ 0.05.

### Reduced transcription of PSI and PSII core genes in Δ*glgC* mutants

Given the enrichment of photosynthesis-related GO terms among downregulated genes, we analyzed TPM values of core photosystem genes in detail. For PSII (Figure 6A, Supplementary Figure S3A), transcripts of *psbA2* (*slr*1311) and *psbA3* (*sll*1867, encoding the D1 protein) accumulated significantly at EoN relative to EoD in all strains, although levels were reduced in the Δ*glgC* mutants at EoN, albeit only significantly for *psbA3*. Expression of *psbB* (*slr*0906) was significantly reduced at EoN in all strains, with lower transcript levels in mutants compared to WT at both time points. Transcript levels of *psbC* (*sll*0851) remained unchanged in the WT, while both Δ*glgC* mutants accumulated significantly fewer transcripts at the EoN. Transcript abundance of *psbD1* (*sll*0849) was basically lower than that of *psbD2* (*slr*0927), and while both homologs were stable in transcript abundance in the WT, they were reduced in both mutants at EoN.

**Figure 6.**
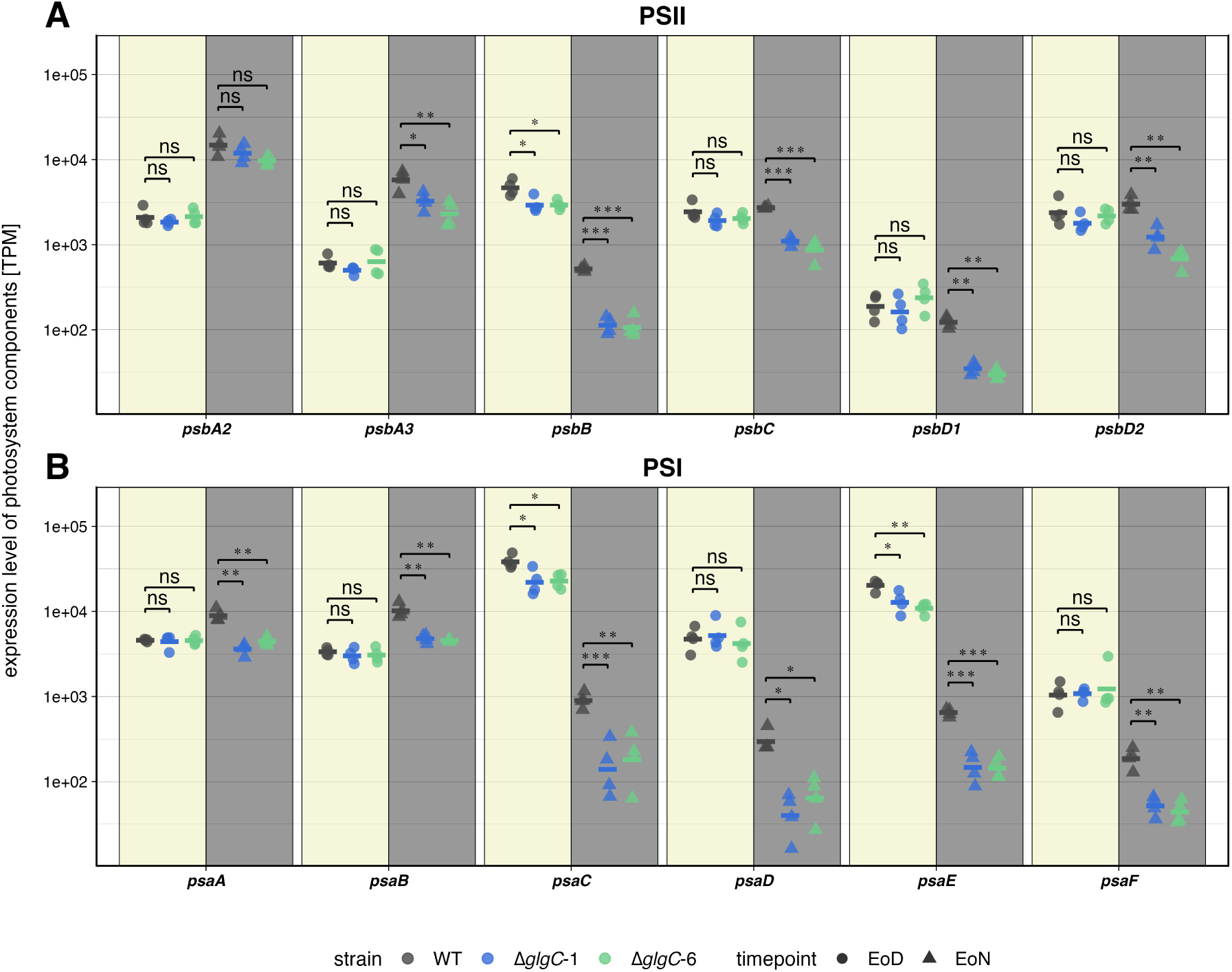
Daytime-specific transcript accumulation of photosynthetic light reaction genes. (A) TPM values of PSII-encoding genes. (B) TPM values of PSI-encoding genes. Data represent the average of four biological replicates. Individual TPM values are provided in Supplementary Data File 1. Asterisks denote statistical significance between genotypes as calculated with the Student’s t-test (*\**: *p ≤ 0.05, \*\**: *p ≤ 0.01, \*\*\**: *p ≤ 0.001*, ns: not significant).

Regarding PSI genes (Figure 6B, Supplementary Figure S3B), *psaA* (*slr*1834) was significantly upregulated at EoN only in the WT. The mutant lines Δ*glgC*-1 and Δ*glgC*-6 accumulated significantly less *psaB* (*slr*1835) transcript at EoN compared to WT. For *psaC* (*ssl*0563), *psaD* (*slr*0737), *psaE* (*ssr*2831), and *psaF* (*sll*0819), all strains showed reduced transcript levels at EoN, with stronger and significant reductions in both Δ*glgC* mutants compared to WT. Additionally, the Δ*glgC* mutants exhibited reduced *psaC* and *psaE* transcript abundance already at EoD.

Overall, Δ*glgC* mutants showed reduced transcript abundance of PSI and PSII core genes, particularly at EoN.

### Transcriptional effects in glycogen-deficient strains are induced by diurnal regimes

To assess whether the observed transcriptional changes in the mutants were driven by the diurnal regime or reflected genotype-specific effects, we performed RNA-seq on samples from batch cultures grown under constant light (100 µmol photons m⁻² s⁻¹) without a dark phase. In comparison to diurnal EoD-samples, all constant-light samples clustered distinctly along PC1, irrespective of genotype (Supplementary Figure S4A). This separation indicates that diurnal rhythmicity, together with glycogen-dependent effects, represents a major determinant of global transcriptional regulation in the Δ*glgC* mutants.

Notably, WT and Δ*glgC* mutant samples did not separate under constant light conditions, suggesting only minor transcriptional differences between genotypes in the absence of a light–dark cycle (Supplementary Dataset S2). Consistently, only a small number of genes were found to be differentially expressed between Δ*glgC* mutants and the WT, 18 DEGs in Δ*glgC*-1 and 233 DEGs in Δ*glgC*-6 (Supplementary Figure S4B, C). GO term analysis of the Δ*glgC*-6 mutant (Supplementary Figure S4D) revealed an enrichment of downregulated genes associated with “photosynthesis,” representing the only overlap with GO terms identified in EoD samples obtained under diurnal conditions (Fig. 5C). Taken together, these results suggest that the observed transcriptional changes in glycogen-deficient strains are largely driven by the diurnal regime rather than by genotype alone.

### Glycogen deficiency leads to severe ATP depletion at end of night

Consistent with the metabolite PCA (Figure 3C), only minor metabolic differences between WT and mutants were observed at EoD, whereas pronounced differences emerged at EoN (Figure 7). At both time points, ADP-glucose levels were strongly reduced in the mutants, confirming the functional disruption of *glgC*. At EoD, phospho*enol*pyruvate (PEP) accumulated approximately 2-fold in the mutants, while most Calvin–Benson–Bassham (CBB) cycle and gluconeogenic intermediates showed non-significant increases.

**Figure 7.**
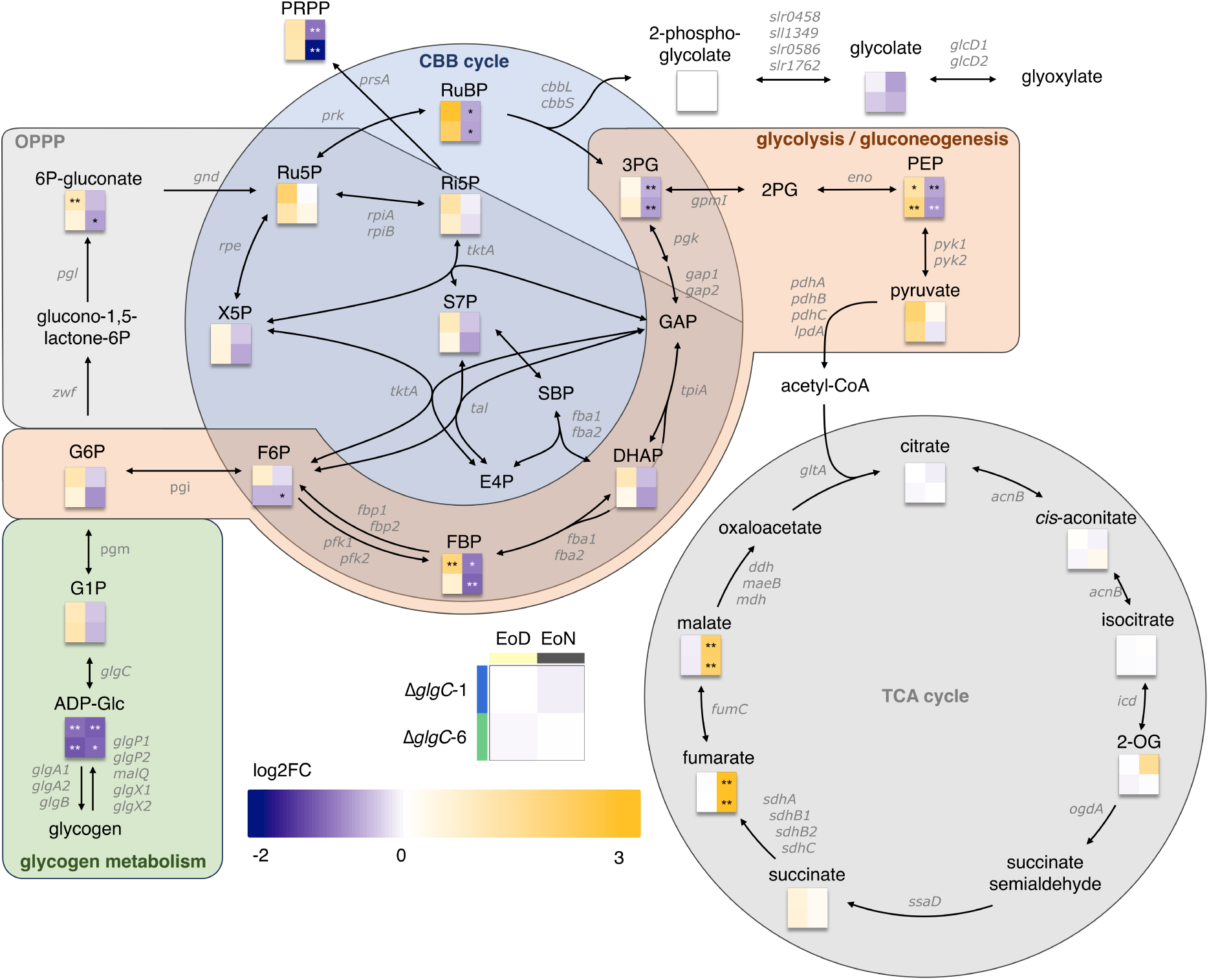
Metabolic response of *ΔglgC* mutants under diurnal cycles. Heatmaps indicate median log₂ fold-changes of metabolite levels in Δ*glgC* mutants relative to WT at EoD and EoN, respectively. Significance thresholds were determined using a two-sided Wilcoxon rank-sum test (*: *p* ≤ 0.05, **: *p* ≤ 0.01, ***: *p* ≤ 0.001). Enzyme-encoding genes are annotated along the arrows. RuBP and 2-phosphoglycolate are indistinguishable in this analysis. Metabolites: ADP-Glc: ADP-glucose; DHAP: dihydroxyacetone phosphate; GAP: glyceraldehyde-3-phosphate; E4P, erythrose-4-phosphate; G1P: glucose 1-phosphate; G6P: glucose 6-phosphate; F6P: fructose-6-phosphate; FBP: fructose-1,6-bisphosphate; 2-PG: 2-phosphoglycerate; 3-PG: 3-phosphoglycerate; 2-OG, 2-oxoglutarate; PEP, phosphoenolpyruvate; PRPP, phospho-ribose-pyrophosphate; RuBP, ribulose-1,5-bisphosphate; Ru5P, ribulose-5-phosphate; SBP, seduheptulose-1,7-bisphosphate; S7P, sedoheptulose-7-phosphate; X5P, xylulose-5-phosphate. Genes encoding relevant enzymes: *acnB*: bifunctional aconitate hydratase / methylisocitrate dehydratase; *cbbL*: RuBisCO large subunit; *cbbS*: RuBisCO small subunit; *ddh*: 2-hydroxyacid dehydrogenase, *eno*: phosphopyruvate hydratase; *fumC*: fumarate hydratase; *fba1*: class I fructose-phosphate aldolase, *fba2*: class II fructose-bisphosphate aldolase; *fbp1*: class II fructose-1,6-/ seduheptuloses-1,7-bisphosphate phosphatase, *fbp2*: class I fructose-1,6-bisphosphate phosphatase; *gap1*: type I glyceraldehyde-3-phosphate dehydrogenase (glycolytic variant); *gap2*: type I glyceraldehyde-3-phosphate dehydrogenase (gluconeogenetic variant); *glcD1D2*: glycolate dehydrogenase 1 & 2; *glgA1* & *glgA2*: glycogen synthase; *glgB*: branching enzyme; *glgC*: adenylyl-glucose pyrophosphorylase; *glgP1* & *glgP2*: glycogen phosphorylase; *glgX1* & *glgX2*: glycogen debranching protein; *gltA*: citrate synthase; *gnd*: phosphogluconate dehydrogenase; *gpmI*: phosphoglycerate mutase; *icd*: isocitrate dehydrogenase; *malQ*: 4-α-glucanotransferase; *maeB*: malic enzyme; *ogdA*: 2-oxoglutarate decarboxylase; *pdhABC* & *lpdA*: pyruvate dehydrogenase E1; *pfk1* & *pfk2*: phosphofructokinase; *pgl*: 6-phosphogluconolactonase; *pgm*: phosphoglucomutase; *prk*: phosphoribulokinase; *pyk1* & *pyk2*: pyruvate kinase, *rpe*: ribulose-phosphate-3-epimerase; *pgi*: glucosephosphate isomerase; *pgk*: phosphoglycerate kinase; *rpiA*: ribose-5-phosphate-isomerase RpiA, *rpiB*: ribose-5-phosphate isomerase RpiB; *sdhAB1B2C*: succinate dehydrogenase / fumarate reductase; *ssaD*: succinate-semialdehyde dehydrogenase; *tal*: transaldolase; *tktA*: transketolase; *tpiA*: triose-phosphate isomerase, *mdh*: malate dehydrogenase; *zwf*: glucose 6-phosphate dehydrogenase; *slr*0458 & *sll*1349 & *slr*1762: 2-phosphoglycolate phosphatase.

At EoN, central carbon metabolites exhibited the opposite trend and were less abundant in the mutants, with significant reductions observed for fructose 1,6-bisphosphate, 3-phosphoglycerate, phospho*enol*pyruvate, and RuBP. Although WT cells also showed reductions at EoN (Supplementary Figure S5), these changes were more pronounced in the mutants, consistent with the absence of glycogen reserves. As RuBP degrades into 2-phosphoglycolate during IC analysis we only get one signal for both compounds. Because 2-phosphoglycolate formation is expected to be negligible in darkness, we interpret the RuBP/2-phosphoglycolate signal primarily as RuBP.

A notable exception was the 4-fold accumulation of malate and fumarate at EoN, observed exclusively in Δ*glgC* mutants. Transcriptomic data supported this response, with increased expression of *sdhB2* (*sll*0823), *fumC* (*slr*0018), and *maeB* (*slr*0721) (Supplementary Figure S6). While *sdhB2* and *fumC* catalyze succinate-to-malate conversion, *maeB* encodes a malic enzyme converting malate to pyruvate with CO₂ release. Upregulation of *maeB* but not *mdh* (*sll*0891) suggests preferential routing through a carbon-losing reaction despite overall carbon depletion.

Given the observed metabolic depletion, we quantified adenylate energy charge (AEC) using ATP, ADP, and AMP concentrations determined by IC-MS with ¹⁵N-labelled internal standards (Figure 8A, B). At EoD, AEC values ranged from 0.5 to 0.55 in all strains. At EoN, WT cells maintained an AEC of 0.64, whereas Δ*glgC*-1 and Δ*glgC*-6 exhibited significantly lower values (0.44 and 0.46, respectively).

**Figure 8.**
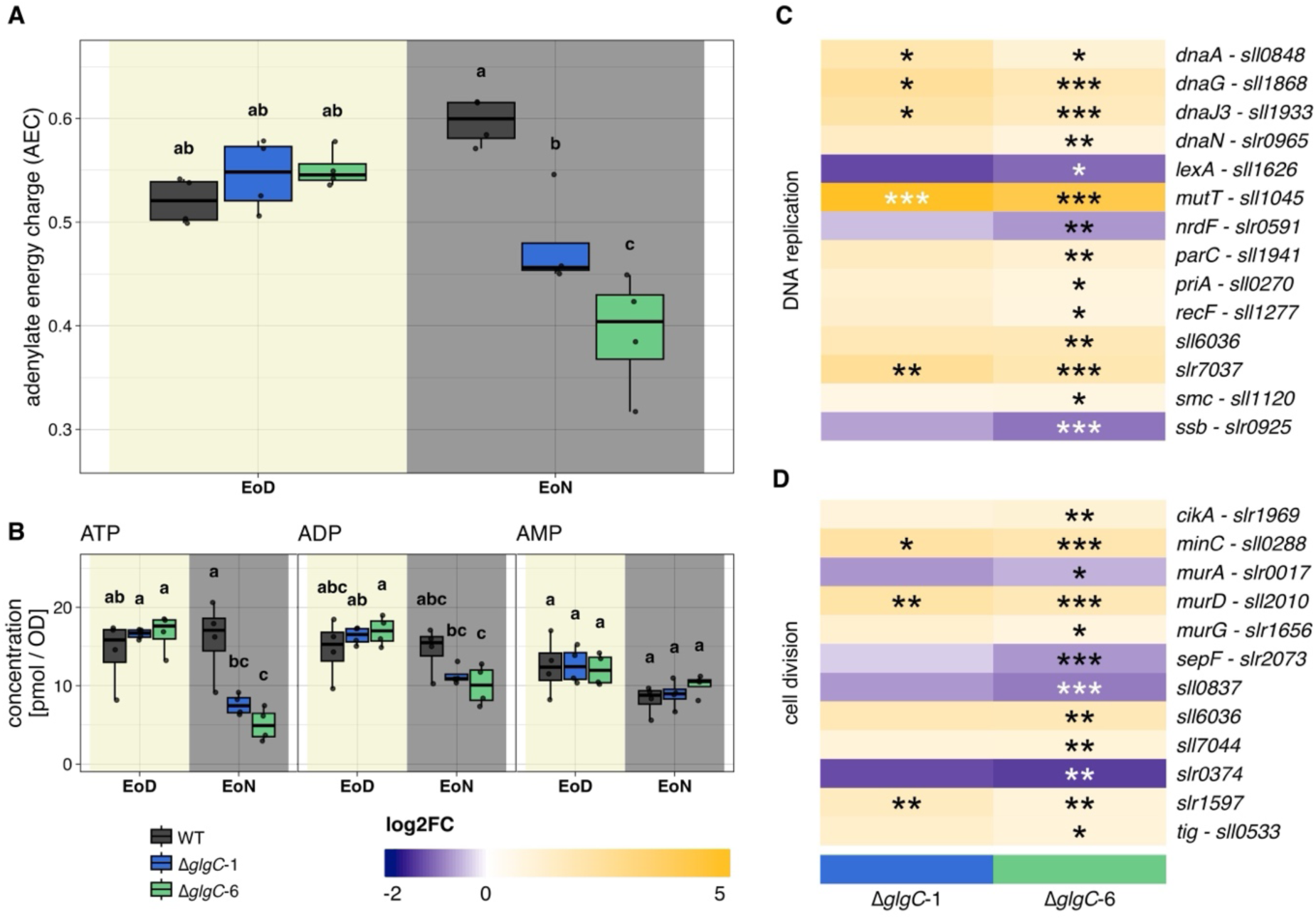
Adenylate energy charge (AEC) and cell division-related transcription in WT and Δ*glgC* mutants. (A) AEC calculated as AEC = ([ATP] + 0.5[ADP])/([ATP] + [ADP] + [AMP]). (B) Concentrations of ATP, ADP, and AMP used to calculate AEC. Letters indicate significance (Kruskal-Wallis test with Tukey post-hoc, *p* ≤ 0.05; n = 4). (C–D) Heatmaps of mean log₂ fold-changes in transcripts associated with DNA replication and cell division at EoN in mutants versus WT (CyanoCyc annotation). Stars denote significance (*: *p* ≤ 0.05, **: *p* ≤ 0.01, ***: *p* ≤ 0.001). Full GO term list is provided in Supplementary Table S1.

To assess whether reduced energy charge affected ATP-dependent processes, we examined transcripts associated with DNA synthesis and cell division (Figure 8C, D). Several genes involved in DNA replication (*dnaA* (*sll*0848), *dnaG* (*sll*1868), *dnaJ3* (*sll*1933)) were significantly upregulated, with additional responses specific to Δ*glgC*-6. Transcripts of *mutT* (*sll*1045), involved in repair of oxidative DNA damage, were strongly increased at both EoN and EoD. Core cell-division responses included increased transcription of *murD* (*sll*2010), *minC* (*sll*0288), and *slr*1597. Additional DEGs in Δ*glgC*-6 included *sepF* (*slr*2073), *sll*6036, and *cikA* (*slr*1969), while *murA* (*slr*0017) was downregulated, suggesting suppression of early cell wall biosynthesis despite downstream activation. Responses were qualitatively and quantitatively similar between mutants, indicating that Δ*glgC*-6 is representative of the genotype.

### Identification of secondary site mutations in Δ*glgC* mutants

Given the observed differences between the Δ*glgC*-1 and Δ*glgC*-6 mutant lines in growth, transcriptional and metabolic response, we decided to perform genome sequencing to identify possible causative secondary site mutations (SSMs). We called 1,942 SSMs that are exclusive to mutants’ genomes (Supplementary Dataset S6). These variants potentially affect 134 genes, with 31 genes carrying high impact variants, that is frame-shift mutations leading to a premature stop codon (Supplementary Fig. S7I-L). Among the affected genes, we identified candidates with high relevance to glycogen and energy metabolism, such as *glgX*2 (*slr*1857), *pgi* (*slr*1345), *ndbC* (*sll*1484), *psaA* (*slr*1834), *petC2* (*slr*1185), or *ccmS* (*slr1*911). Importantly, the variant allele frequencies for the SSMs was mostly below <50% (Supplementary Figure S7B-D), indicating that the polyploid genomes were not fully segregated for the mutations and can thus not be equated to a full knockout. However, partial segregation may still affect protein levels as a result of gene dosage (Nagy et al., 2021).

## DISCUSSION

Glycogen-deficient *Synechocystis* strains have been characterized previously. However, the physiological and regulatory strategies by which these mutants cope with the absence of a daytime carbon sink and the lack of carbon reserves during the night remain poorly understood. Here, we provide mechanistic insight into how *Synechocystis* Δ*glgC* mutants initially respond to the loss of its primary carbon and energy buffer under diurnal conditions, linking glycogen deficiency to acceptor limitation during the day and an ATP crisis during the night. While the mutant cells get growth arrested after very few diurnal cycles with 12-h light/dark regime, we here took a snapshot into the initial response towards glycogen deficiency. Importantly, the microfluidic and batch cultivation datasets presented here provide complementary, but not directly comparable, insights into the physiological consequences of glycogen deficiency. The two systems were designed to address different levels of resolution, with microfluidics capturing single-cell growth dynamics and batch cultures enabling transcriptomic and metabolomic analyses under defined diurnal conditions. A conceptual summary of our findings is provided in Figure 9.

**Figure 9.**
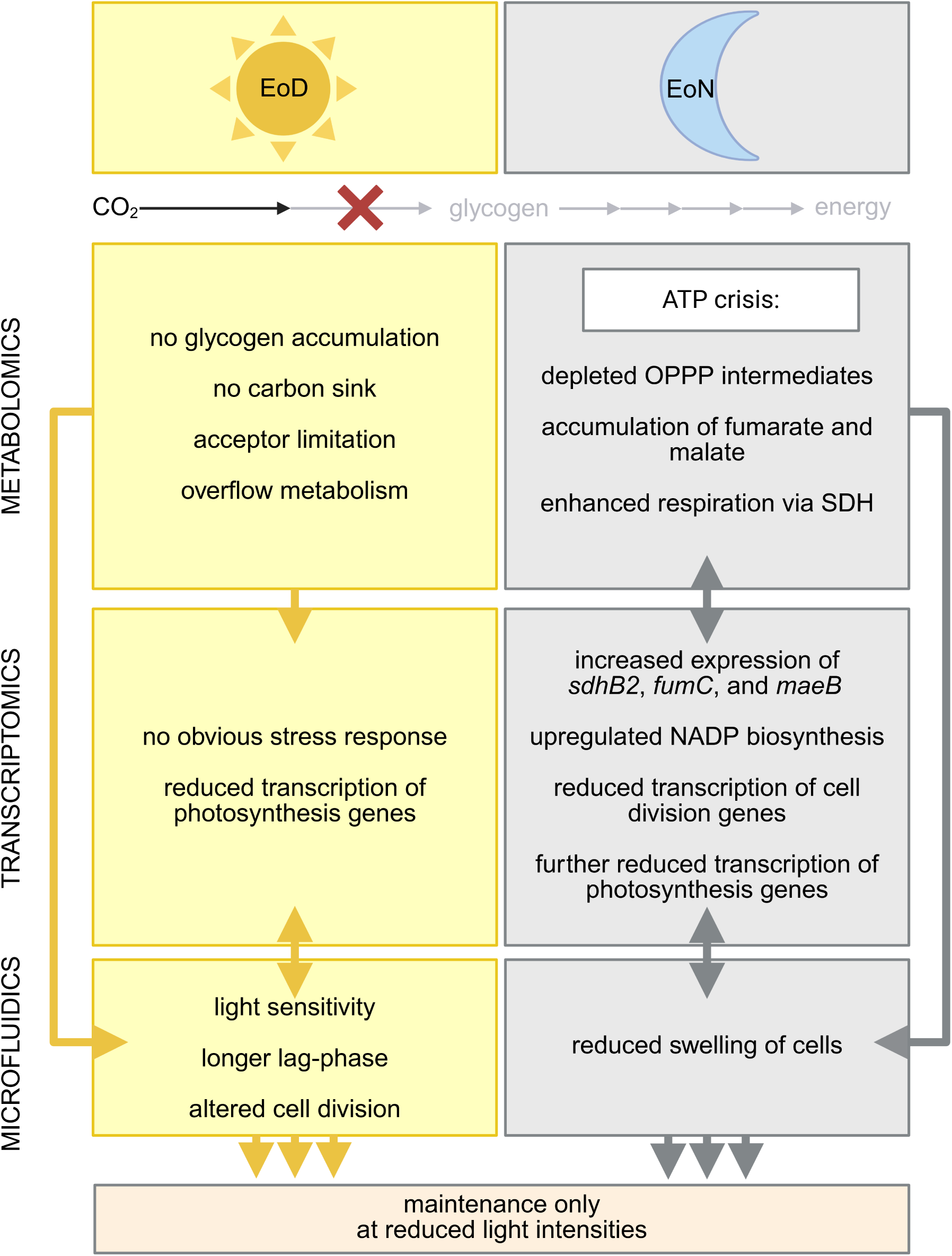
Model of glycogen-dependent metabolic regulation in *Synechocystis* under diurnal cycles. Lack of glycogen alters transcriptomic, metabolomic, and phenotypic states. During the day, photosynthesis-related genes are downregulated, likely to mitigate acceptor limitation associated with the missing carbon sink, while overflow metabolism may help maintain redox balance. At night, absence of glycogen results in an ATP crisis. Carbon is redirected via succinate dehydrogenase toward fumarate and malate (upregulation of *sdhB2*, *fumC*, *maeB*), supporting ATP synthesis. Genes involved in cell division and cell wall synthesis are also affected. Redox balance may also be affected as NAD(P) biosynthesis and other redox-responsive pathways are regulated at transcriptional level. Together with depletion of OPPP intermediates and reduced photosynthesis transcripts, this leads to prolonged lag phases and delayed resumption of photosynthetic activity at dawn.

### Δ*glgC* mutants reduce photosynthetic capacity to mitigate acceptor limitation

Single-cell cultivation in the recently developed microfluidic platform (Witting et al., 2025) under precisely controlled light intensity (Figure 2A, B) revealed physiological constraints that were previously masked in bulk cultures. Consistent with earlier reports (Miao et al., 2003; Gründel et al., 2012; Hickman et al., 2013), Δ*glgC* mutants displayed a pronounced photosensitive phenotype. The defined illumination regime enabled us to determine that Δ*glgC* cells lose viability at light intensities exceeding 35 µmol photons m^−2^ s^−1^, whereas the WT reached maximal growth at approximately 80 µmol photons m^−2^ s^−1^ (Figure 2A, B). These intensities are lower than those commonly applied in shake-flask cultures, where cellular self-shading inevitably leads to light gradients and transient light exposure. In contrast, cells in the microfluidic chip grow as a monolayer and are continuously exposed to the applied light intensity.

Notably, the margin between optimal growth and light-induced lethality in Δ*glgC* mutants was remarkably narrow (≈4 µmol photons m^−2^ s^−1^ for both mutants), highlighting the limited physiological flexibility of cells lacking a carbon storage buffer (Sun et al., 1999). Besides metabolite overflow as a valve (Cano et al., 2018), this photosensitivity is likely linked to an insufficient regeneration of the electron acceptor NADP⁺ as a consequence of the missing glycogen sink. During illumination, linear electron flow produces ATP and NADPH at a ratio of ∼1.285, whereas carbon fixation via the CBB cycle requires a ratio of ∼1.5. The additional ATP demand is typically met via cyclic electron flow around PSI, which does not generate NADPH (Kramer and Evans, 2011).

While photosensitive phenotypes were observed in the microfluidic system (Figure 2A, B), they are consistent with the transcriptional downregulation of photosynthesis-related genes observed in batch cultures, suggesting a common response to redox imbalance despite differences in experimental setup. In contrast to what might be expected under photo-oxidative stress, GO-term enrichment analysis did not reveal a general upregulation of stress-response or photoprotective pathways (Figure 5). Instead, only a limited number of individual stress-related genes were differentially expressed (Supplementary Figure S8). Rather, a consistent downregulation of photosynthesis-related genes was observed, including reduced transcript levels of PSI and PSII components and enrichment of GO terms such as “photosynthesis” and “photosynthesis, light reaction” among downregulated genes at both EoD and EoN (Figures 5, 6). This transcriptional pattern suggests an active reduction of photosynthetic capacity as a strategy to alleviate acceptor limitation caused by excess reducing equivalents. Though changes on transcript level do not necessarily translate into physiological changes, in this case, comparable changes were determined earlier in physiological experiments. In Δ*glgC* mutants, linear (Miao et al., 2003; Holland et al., 2016) and cyclic electron flow (Holland et al., 2016; Cano et a., 2018) were found impaired and the NADPH pool highly reduced (Holland et al., 2016).

Photosynthetic capacity in *Synechocystis* is tightly coupled to its ability to channel fixed carbon into metabolic sinks (Grund et al., 2022; Muth-Pawlak et al., 2024). Consequently, glycogen deficiency necessitates compensatory mechanisms to prevent over-reduction of the photosynthetic electron transport chain, as NADPH oxidation to NADP⁺ becomes limiting (Jackson et al., 2015; Holland et al., 2016; Cano et al., 2018). Overflow metabolism can act as an alternative outlet by dissipating ATP and NADPH through secretion of organic acids under nitrogen limitation (Carrieri et al., 2012) or glutamate under nitrogen-replete conditions (Kato et al., 2024; Forchhammer et al., 2025). Because overflow metabolites were not directly quantified here, we cannot conclusively determine whether photosynthetic downregulation precedes or follows overflow metabolism.

### Glycogen is essential for dawn recovery and diurnal energy homeostasis

Photoautotrophic *Synechocystis* does not grow in complete darkness (Shinde et al., 2020), although glucose-tolerant strains can proliferate under light-assisted heterotrophy (LAH) with minimal daily illumination (Anderson and McIntosh, 1991). Reduced growth of glycogen-deficient mutants under diurnal regimes has been reported previously (Gründel et al., 2012; Shimakawa et al., 2014; Cano et al., 2018; Shinde et al., 2020, reviewed in Welkie et al., 2020), but these studies relied primarily on OD measurements. By contrast, the microfluidic platform enabled us to disentangle cell-size increases from cell-division events, providing a more resolved view of growth dynamics (Figure 2).

Our analyses revealed that neither WT nor Δ*glgC* strains divide during the night and that growth impairment in Δ*glgC* mutants primarily results from an extended lag phase at dawn. This phenotype coincided with the absence of the nocturnal cell-swelling observed in WT cells (Figure 2). Metabolomic profiling indicated depletion of oxidative pentose phosphate pathway (OPPP) intermediates at EoN in glycogen-deficient strains (Figure 7). These metabolites are essential for priming photosynthesis and carbon fixation at dawn by ensuring appropriate NADPH levels and precursor availability for the CBB cycle (Shimakawa et al., 2014; Shinde et al., 2020; Tanaka et al., 2023). Concurrently, transcripts encoding photosystem components remained downregulated, further compromising the capacity to resume photoautotrophic growth (Figures 5F, 6).

Without sufficient metabolic priming of the CBB cycle and adequate expression of photosynthetic machinery, Δ*glgC* cells are unable to efficiently fix CO₂ at dawn, resulting in delayed heterotrophy-to-autotrophy transition. The progressive lengthening of the lag phase with increasing dark duration suggests that transient carbon reserves may partially compensate for glycogen deficiency over short periods, whereas extended darkness leads to pronounced starvation. The loss of synchronized cell divisions in Δ*glgC* mutants, in contrast to the tightly coordinated divisions in the WT at dawn (Figure 2), further supports this interpretation. Together with the cell-swelling phenotype in the WT, these observations suggest that glycogen degradation fuels preparatory processes for cell division, a capability that is lost in Δ*glgC* mutants.

### Glycogen deficiency induces an ATP crisis during the night and delays cell division

Consistent with the observed starvation phenotype, metabolomic analyses indicated that Δ*glgC* mutants are unable to maintain sufficient cellular energy levels throughout the night. Specifically, we observed a marked reduction in adenylate energy charge (AEC) at EoN (Figure 8), reflecting impaired ATP synthesis due to the absence of glycogen-driven respiration. Stable ATP levels are critical for ATP-demanding processes, such as chromosome replication, cell wall biosynthesis, and division machinery assembly. Large fluctuations in intracellular ATP have been linked to reduced growth rates in *E. coli* (Lin and Jacobs-Wagner, 2022). The relatively constant ATP levels in the WT contrast sharply with the declining AEC in Δ*glgC* mutants, supporting a causal link between energy limitation and impaired growth recovery. The AEC values measured here are comparable to those reported after circadian disruption in *Synechococcus* following a dark pulse at subjective dawn, where Δ*glgC* strains showed increased sensitivity (Pattanayak et al., 2014).

Given the absence of nocturnal cell swelling phenotype in Δ*glgC* cells, we examined transcriptional responses of the cell-division machinery (Figure 8D). Many division-related proteins bind ATP (or GTP in the case of FtsZ), and chromosome replication, divisome assembly, and cell-wall synthesis are highly energy-demanding processes (Sekimizu et al., 1987; Rueda et al., 2003; Barrows and Goley, 2021). At EoN, Δ*glgC* mutants showed increased transcript levels of *minC*, *murD*, and *slr*1597. MinC is a key regulator of division-site placement in *Synechocystis* (Mazouni et al., 2004) and, as in *E. coli*, can inhibit FtsZ-ring formation when overexpressed (de Boer et al., 1990; Pichoff and Lutkenhaus, 2001). Upregulation of MurD and MurG suggests enhanced cell-wall biosynthetic potential, whereas reduced *murA* expression may indicate negative regulation at the pathway’s committed step (Bhagavan and Ha, 2015). Accumulation of GlcNAc supports this interpretation (Supplementary Figures S5, S9). Additional DEGs, including *sepF*, which stabilizes FtsZ rings, further point to altered regulation of cell division.

Because cell division in *Synechococcus* is under circadian control (Mori et al., 1996) and clock amplitude depends on cellular energy status (Phong et al., 2013), energy depletion may indirectly perturb cell-cycle timing via effects on the circadian oscillator, providing an additional layer of interaction between ATP homeostasis and division control.

The metabolic and transcriptional findings are supported by the reduced growth recovery and extended lag phases observed at the single-cell level, although a direct quantitative comparison between systems is not possible.

Together, these findings mechanistically link ATP depletion to delayed cell division.

### Compensatory metabolic rerouting partially restores energy and redox balance

One of the most prominent metabolic signatures of Δ*glgC* mutants at night was the accumulation of the TCA cycle intermediates fumarate and malate (Figure 7), the only metabolites detected by IC–MS that increased during darkness (Supplementary Dataset S3). This accumulation was accompanied by elevated transcript levels of *sdhB2, fumC*, and *maeB* in Δ*glgC*-6 (Supplementary Figure S6). The malic enzyme MaeB catalyzes the conversion of malate to pyruvate and plays a central role in directing malate flux (Bricker et al., 2004; Katayama et al., 2022). Because pyruvate levels remained unchanged, a primary carbon-balancing function appears unlikely. Notably both enzymes depend on NAD^+^ as a cofactor.

While proteolysis and subsequent breakdown of amino acids could lead to metabolic imbalances and the observed malate accumulation, we observe no evidence of increased amino acid degradation at EoN at transcriptomic level. Malate may also be produced by the activity of phosphoenolpyruvate carboxylase (PepC), however transcription of the gene is not significantly increased, and fumarate and malate accumulation have previously been shown to inhibit its enzymatic activity (Takeya et al., 2017). Instead, we propose that Δ*glgC* mutants favor continued activity of the succinate dehydrogenase (SDH) complex, supported by upregulation of SDH, FumC, and MaeB. SDH reduces plastoquinone during succinate oxidation (Cooley and Vermaas, 2001), thereby feeding electrons into the shared photosynthetic–respiratory electron transport chain (Battchikova et al., 2011; Lea-Smith et al., 2016). As SDH has been suggested as a major contributor to plastoquinone reduction in *Synechocystis* (Cooley et al., 2000), enhanced SDH activity could sustain residual respiratory electron flow and ATP synthesis during the night. Nonetheless, reduced expression of cytochrome b_6_f, CydAB, and COX (Supplementary Figure S10) suggests that overall respiratory capacity remains lower than in the WT, consistent with the reduced AEC.

Finally, we observed significant upregulation of type II NAD(P)H dehydrogenase genes *ndbA* and *ndbC*. NdbA is associated with LAH growth and maintenance of photosystem integrity (Huokko et al., 2019), whereas NdbC has been implicated in carbon partitioning, sugar catabolism, and cell division (Huokko et al., 2017). Because *ndbC* deletion leads to glycogen accumulation, its overexpression in Δ*glgC* mutants may represent an attempt to reroute carbon flux in the absence of glycogen. Together with broad transcriptional changes in redox-related pathways (Figure 5, Supplementary Dataset S1), this points to a pronounced dysregulation of NAD(P)(H) homeostasis. Using genome sequencing, we detected a high-impact frame-shift SSM in *ndbC* with an allele frequency of 30% in the genome of Δ*glgC*-6 (Supplementary Figure S7, Supplementary Dataset S6). Based on the gene-dosage effect (Birchler & Veitia, 2012; Nagy et al., 2021)), this rather low allele frequency may still impact the protein abundance of NdbC and thus reduce the efficiency of the compensatory strategy. We hypothesize that this SSM likely contributes to the observed stronger phenotype in comparison to Δ*glgC*-1.

In line with this interpretation, genes involved in *de novo* NAD(P) biosynthesis and salvage pathways (*nadA–C*, *nadM*, *nadV*) were upregulated at EoN, along with *nadk1*, which encodes an NAD kinase involved in balancing NAD⁺/NADP⁺ ratios (Ishikawa et al., 2021). Shifting the redox balance towards NADP⁺ may represent a conserved strategy to ensure sufficient electron acceptor availability at dawn, as previously observed in *Synechococcus* (Shinde et al., 2020) and appears to be maintained despite the loss of glycogen synthesis. However, this adaptation may compound malate and fumarate accumulation by decreasing the availability of NAD^+^, the co-substrate of malate metabolizing enzymes Mdh, Ddh and MaeB.

## Conclusion

In summary, while the experimental setups differ, both approaches consistently demonstrate that glycogen functions as a central metabolic integrator in *Synechocystis*, stabilizing both redox balance during the day and sustaining cellular energy supply during the night. In the absence of this carbon store, Δ*glgC* mutants experience acceptor limitation under illumination, triggering downregulation of photosynthetic gene expression to prevent over-reduction. During the dark phase, depletion of glycogen-derived substrates results in a pronounced ATP deficit that compromised metabolic priming for the subsequent light period, delaying the reestablishment of photoautotrophic growth and disrupting the timing of cell division. Glycogen-deficient cells partially compensate for these constraints by extensive metabolic and transcriptional reprogramming, including maintained flux through SDH, as well as upregulation of NAD(P) biosynthesis and type II NAD(P)H dehydrogenases. Though a fully coordinated compensatory response is missing, cells remain resilient, surviving at least 12 h of darkness. However, cells run into growth arrest after just very few diurnal cycles. Together, these observations establish a mechanistic framework linking glycogen metabolism to diurnal fitness through coordinated control of carbon storage, energy homeostasis, and transcriptional regulation. While integration of single-cell and omics data within a unified experimental platform would further strengthen these conclusions, the combined use of complementary approaches already provides a robust framework to understand the systemic consequences of glycogen deficiency.

Beyond physiological implications, these results are consistent with evolutionary models of cyanobacterial endosymbiosis in which the loss of autonomous carbon storage constrains free-living growth while promoting metabolic dependence on a host. In such a context, host-mediated energy provisioning during dark periods could stabilize endosymbiont populations and enable tight metabolic integration. Our data therefore underscores metabolic connectivity as a fundamental principle of shaping host control over endosymbiont population dynamics.

## MATERIAL & METHODS

### Standard cultivation conditions

*Synechocystis* sp. PCC 6803 substr. Kazusa wild type (WT) was obtained from the University of Rostock (M. Hagemann). Cultures were propagated in shake flasks in an Infors HT shaker using BG11 medium (Rippka et al., 1979) under constant illumination (30 µmol photons m^−2^ s^−1^) at 30 °C and 110 rpm in unbaffled flasks filled to 20% of the nominal volume. For Δ*glgC* mutants, selective pressure was maintained by supplementing the medium with kanamycin (50 µg mL^−1^). In experimental cultures, Kanamycin was omitted.

### Generation of glycogen-deficient mutants

Δ*glgC* knockout mutants were generated by insertional inactivation of the *glgC* gene via homologous recombination using a kanamycin resistance cassette. A schematic overview of the mutagenesis strategy is shown in Supplementary Figure S11. The *glgC* (*slr*1176) coding sequence was amplified from genomic DNA by PCR using primers JS69 (GGCATCAACGGCGTTGGAAA) and JS70 (GGCACCACTTCCACCGACTG) and cloned into pJET1.2 (Thermo Scientific). The resulting construct was digested with PsiI, and a kanamycin resistance cassette excised from pUC4K by HincII digestion was ligated into the PsiI site within the *glgC* open reading frame (Berwanger et al., 2023). The final construct (pJS22) was verified by sequencing. Natural transformation of *Synechocystis* was performed as described previously (Ermakova et al., 1993). Transformants were selected on BG11 agar plates containing 25 µg mL^−1^ kanamycin and subsequently transferred to plates containing 50 µg mL^−1^ and 100 µg mL^−1^ kanamycin to promote full segregation. Complete replacement of the WT allele was confirmed by PCR using primers JS53 (GGCCGGAAAGTATCGCCT) and JS54 (AAACAATCGCAGGCCGCC). Two fully segregated mutant lines derived from independent transformation events (Δ*glgC*-1 and Δ*glgC*-6) were selected for further experiments.

### Determination of glycogen content

Intracellular glycogen content was determined based on the method described by Gründel et al. (2012), with modifications according to Makowka et al. (2020) and Lucius et al. (2021). 2 mL cells with an OD_750_ of 0.8 were harvested after 4 d cultivation in a Multi-Cultivator 1000-OD photobioreactor at 30 °C and 100 µmol photons m^−2^ s^−1^ continuous light and frozen at −20 °C. Cell pellets were thawed on ice and washed three times with 1 mL double-distilled water, followed by centrifugation at 10,000 x *g* for 10 min. Pellets were resuspended in 400 µL 30% (w/v) KOH and incubated at 95 °C for 2 h. Glycogen was precipitated by addition of 1.2 mL ice-cold 96% ethanol and incubation at −20 °C overnight.

Samples were centrifuged at 10,000 x *g* for 10 min at 4 °C, washed once with 70% ethanol and once with 98% ethanol, and air-dried at 30 °C to remove residual ethanol. Pellets were resuspended in 1 mL sodium acetate buffer (100 mM, pH 4.5) and incubated with 100 µL amyloglucosidase (16.8 U mL^−1^; Sigma-Aldrich/Roche) at 60 °C for at least 2 h to hydrolyze glycogen to glucose. Samples were centrifuged at 10,000 x *g* for 10 min, and glucose concentrations were quantified colorimetrically using *o*-toluidine reagent. Absorbance was measured at 635 nm using an Infinite M Nano plate reader (Tecan).

### Cultivation in a microfluidic chemostat

Microfluidic cultivation of WT and Δ*glgC* mutants was performed using a previously described platform (Witting et al., 2025). Cells were grown in monolayers within growth chambers measuring 60 x 60 x 1 µm and monitored by time-lapse microscopy using an inverted microscope (Ti-E, Nikon, Japan) equipped for phototrophic growth. BG11 medium was continuously supplied at 200 nL min^−1^ using a syringe pump (Nemesys, Cetoni, Germany), and cultivation was performed at 30 °C.

Chambers were inoculated with exponentially growing cells cultivated in a photobioreactor (Multi-Cultivator 1000-OD, Photon Systems Instruments, Czech Republic) D uring time-lapse imaging phase-contrast and chlorophyll fluorescence images (excitation 514/30 nm, dichroic mirror 561 nm, emission 629/56 nm) were captured.

Two experimental regimes were applied: (i) diurnal experiments with homogenous illumination of the entire chip at 30 µmol photons m^−2^ s^−1^, and (ii) parallel cultivation under constant light at multiple light intensities to determine light-dependent growth rates.

The resulting image sequences were sorted into three categories by visually inspecting the data: (i) observable growth and cell division of the inoculated cell; (ii) observable growth and cell division of the inoculated cell, however determination of growth rates not possible (e.g. due to poor image quality); (iii) no growth or cell division of the initial cell observed. Images of category (ii) were not further evaluated. Image stacks of category (i) were pre-processed in Fiji to correct for stage drift and crop chamber boundaries. Cell segmentation and downstream analyses were performed as described previously (Witting et al., 2025) using custom Python scripts. Growth rates were calculated by linear regression of ln-transformed colony area over time.

### Cultivation for combined diurnal transcriptomic and metabolomic analyses

Pre-cultures (100 mL) of WT, Δ*glgC*-1, and Δ*glgC*-6 were grown under standard cultivation conditions, including Kanamycin. After 11 d, cultures were harvested by centrifugation (2 × 10 min at 3,000 x *g*), washed once with fresh BG11 medium, and resuspended in 20 mL BG11 without Kanamycin. Main cultures were inoculated to an initial OD₇₅₀ of 0.5 in a final volume of 80 mL without Kanamycin.

Cultivation was performed in a Multi-Cultivator 1000-OD photobioreactor (Photon Systems Instruments) at 30 °C and 100 µmol photons m^−2^ s^−1^. Cultures were acclimated for 24 h under continuous light, followed by a 12 h dark/12 h light cycle starting with a dark phase to resynchronize the circadian clock. End-of-day (EoD) samples were collected after 11 h of illumination (47 h after inoculation), and end-of-night (EoN) samples after 11 h of darkness (59 h). The experiment was terminated after 60 h.

### Sampling for transcriptomic and metabolomic analyses

At each sampling time point, aliquots were collected for metabolite extraction, RNA isolation, and optical density measurements. For metabolite analysis, 5 mL of culture were rapidly transferred to ice-cold 0.9% NaCl in a vacuum filtration apparatus (Merck Millipore) and filtered onto nylon membranes (25 mm diameter, 0.45 µm pore size). Filters were immediately transferred to 2 mL tubes containing steel and glass beads (Retsch) and flash-frozen in liquid nitrogen. Total processing time did not exceed 45 s. Sampling during light phases was performed under illumination matching cultivation conditions, whereas dark-phase sampling was conducted under green light.

For RNA isolation, 5 mL of culture were centrifuged at 3,000 x *g* for 10 min at 4 °C, and pellets were flash-frozen in liquid nitrogen. Optical density was measured at 750 nm after 1:10 dilution in 0.9% NaCl using a SpectroStar Nano plate reader (BMG Labtech).

### RNA extraction and sequencing

RNA extraction was performed with the RNeasy® Plant Mini Kit (Qiagen) according to the manufacturer’s instructions. To prepare the RLT buffer, for every 1 mL of RLT buffer 10 µL of 2-mercaptoethanol was added. After thawing the samples on ice for more than 1 hour, 500 µL of RLT buffer was used to resuspend the cell pellets of each sample. The cells were mechanically lysed using 0.5 mm beads (Roth) in the Precellys Evolution cell lyser (Bertin Technologies) which was cooled down to between −5 and −10 °C via the Cryolys (Bertin Technologies, 2013 version) that has been supplied with liquid nitrogen. The lysate was transferred onto the QIAshredder columns and the instructions in the manual were followed. After the washing step with the RWE buffer, 80 µL of a 340 Kunitz/ml DNaseI in RDD buffer (Qiagen RNAse-Free DNase Set) solution was pipetted onto the membrane of the spin column. The DNaseI digest was left to incubate for 30 min at RT. To elute the RNA, 35 µL of RNase-free water was added to the column and centrifuged at 8,000 x *g* for 1 min. The RNA was stored until further use at −80 °C. To prepare the RNA for sequencing, rRNA depletion and library preparation were performed with the Pan-Bacteria riboPOOL kit (siTOOLs) and the bacteria Illumina TruSeq stranded mRNA UDI kit. The sequencing was performed on a NextSeq 2000 (Illumina) using a NextSeq2000 P2 flow cell for 100 cycles and the XLEAP chemistry in single-end mode. The *Synechocystis* sp. PCC 6803 genome assembly ASM972v1 (RefSeq accession no. GCF_000009725.1) and its corresponding gene annotation (RefSeq release v2022-10-16) were used as the basis for all computational analyses. All programs were run with default parameters unless otherwise specified.

### Functional annotation of transcripts

To create a comprehensive functional annotation table, protein domains were identified with InterProScan v5.62-94.0 (Jones et al., 2014) using the databases Pfam v35.0 (Finn et al., 2016), SUPERFAMILY v1.75 (Wilson et al., 2009), and FunFam v4.3.0 (Scheibenreif et al., 2019). A blastp search was run to identify the best hits against the Arabidopsis thaliana TAIR10 gene annotation, with the parameter -max_hsps 1 (NCBI-BLAST+ v2.14.0; Camacho et al., 2009). Functional categories were further assigned using the web application of the eggNOG-mapper v2.1.12 (Cantalapiedra et al., 2021). All obtained information, together with the data from the GFF file and the CyanoCyc database v29.0 (Moore et al., 2024), was integrated into a comprehensive functional annotation table (Supplementary Dataset S1).

### Differential gene expression analysis

The single-end RNA-Seq reads were trimmed using Trimmomatic v0.40 (Bolger et al., 2014) with the TruSeq3-SE.fa adapter file and the parameters MINLEN:60, LEADING:34, TRAILING:34, SLIDINGWINDOW:4:15, and ILLUMINACLIP:2:34:15. Quality of raw and trimmed reads was verified with FastQC v0.12.1 (Andrews, 2010).

For transcripts per million (TPM) calculation, transcript quantification was performed with Kallisto v0.51.1 (Bray et al., 2016) using the quant function, the trimmed reads, the coding sequences and the parameters -l 200 and -s 50.

Following analyses were done in R v4.5.2 (R Core Team, 2025), and plots were created with ggplot2 v3.5.2 (Wickham, 2016). A principal component analysis (PCA) was computed on the mean TPM values per biological replicate with R’s function prcomp and parameter scale=T. Differential gene expression (DGE) analysis was performed with edgeR v4.6.3 (Chen et al., 2025). Technical replicates were summed to obtain counts per biological replicate prior to analysis, and genes with zero counts across all samples were excluded. Genes with a *q*-value below 0.01 were considered significantly differentially expressed.

### Gene Ontology enrichment analysis

Gene Ontology (GO) enrichment analysis was performed using the R package topGO (Alexa and Rahnenfuhrer, 2025). GO annotations were taken from the functional annotation table described above (Supplementary Dataset S1). Enrichment analysis for the ontology biological process (BP) was conducted for the set of up- and downregulated genes separately. For testing, Fisher’s exact test and the classic algorithm of topGO were selected. Only GO terms associated with at least ten genes (parameter nodeSize = 10) were considered. GO terms were semantically clustered using GO-Figure! v1.0.2 (Reijnders and Waterhouse, 2021) with the parameter --max_pvalue 0.05.

### Metabolite analysis using IC–MS

Metabolite analysis was performed following a protocol based on Rabinowitz and Kimball (2007). Briefly, 1 mL of extraction solvent (40:40:20 acetonitrile:methanol:0.1 M formic acid, v/v) containing ribitol (5 µM), dimethylphenylalanine (5 µM) and thio-ATP (2 µM) as internal standards was added to reaction tubes containing nylon filters with harvested cells, as described above. Samples were disrupted in a bead mill (Retsch) at 30 Hz for 2 min using pre-cooled holders to detach cells from the filter and ensure efficient lysis. Subsequently, 90 µL of 15% (w/v) NH_4_OH was added to neutralize the extract.

Nylon filters and steel beads were removed, and samples were incubated at −20 °C for 30 min to precipitate proteins. After centrifugation at 16,000 x *g* for 10 min at 4 °C, the supernatant was transferred to a 15 mL-tube and diluted with 5 mL of ice-cold H₂O. Samples were frozen at −80 °C, lyophilized, and stored at −80 °C until further processing. Prior to analysis, dried extracts were resuspended in 500 µL H_2_O.

Metabolites were analysed using a Dionex ICS-6000 HPIC system coupled to a Thermo Scientific Q Exactive Plus quadrupole–Orbitrap mass spectrometer, as described by Schwaiger et al. (2017). A 10 µL aliquot of the extracted sample was injected into the IC system using a Dionex AS-AP autosampler operated in push partial mode.

Anion exchange chromatography was performed using a Dionex IonPac AS11-HC analytical column (2 mm × 250 mm, 4 µm particle size; Thermo Fisher Scientific) equipped with a Dionex IonPac AG11-HC guard column (2 mm × 50 mm, 4 µm; Thermo Fisher Scientific) and operated at a column temperature of 30 °C. The mobile phase was generated using an eluent generator with a potassium hydroxide cartridge, producing a potassium hydroxide gradient. The flow rate was set to 380 µL min⁻¹, starting at a KOH concentration of 5 mM for 1 min, followed by a linear increase to 85 mM over 35 min, which was held for 5 min. The concentration was then immediately reduced to 5 mM, and the system was equilibrated for 10 min.

Spray stability of aqueous solution was improved using a makeup flow of methanol containing 10 mM acetic acid, delivered by an AXP pump with a flow rate of 150 µL min⁻¹. Electrospray ionisation was performed in the ESI source using the following parameters: sheath gas 30, auxiliary gas 15, sweep gas 0, spray voltage −2.8 kV, capillary temperature 300 °C, S-Lens RF level 45, and auxiliary gas heater temperature 380 °C.

Full-scan spectra were acquired over an m/z range of 60–800 at a resolution of 140,000, with an automatic gain control (AGC) target of 1 × 10⁶ ions and a maximum injection time of 500 ms.

Targeted data evaluation was performed using Skyline (version 25.1, MacCos Lab Software, University of Washington). Peak areas were normalized to internal standards and to OD_750_ values measured at the time of sampling. RuBP dissociates on the anion-exchange column due to the high pH during the chromatographic run and its interaction with the positively charged stationary phase. This dissociation generates 2-phosphoglycolate, which accounts for the major proportion of the measured RuBP/2-phosphoglycolate signal in biological samples. Therefore, we use the RuBP/2-phosphoglycolate signal here primarily as a proxy for RuBP and as a metabolic marker of photosynthetic activity. Statistical analyses were conducted using a custom R script.

## Accession numbers/Data availability

The diurnal RNA-seq data set is available at the European Nucleotide Archive (ENA) with the Accession ID PRJEB111315. The constant-light RNA-seq data is available with the Accession ID PRJEB111882.

The genome assemblies, gene annotation, and raw reads are available in GenBank/ENA under BioProject accession number PRJEB112054.

The microfluidic images and metabolomic data are available at https://git.nfdi4plants.org/schulze.tim1995/Synechocystis_glycogen_knockout_RNA_seq_metabolomics (Add DOI once published).

## ACKNOWLEDGEMENTS

We thank Jeannine Volke for kindly providing the Δ*glgC* mutants. We thank Tobias Busche and his team of the Omics Core Facility NGS Unit at Bielefeld University Medical School OWL and CeBiTec for sequencing of RNA samples. We acknowledge support by the BMBF-funded de.NBI Cloud within the German Network for Bioinformatics Infrastructure, and the CeBiTec compute cluster for computational resources.

## FUNDING

This work was funded by the Deutsche Forschungsgemeinschaft (DFG, German Research Foundation) – SFB1535 - Project ID 458090666.

## AUTHOR CONTRIBUTIONS

JMH and TS performed the diurnal transcriptomics/metabolomics experiment. JMH performed the metabolite extraction and data analysis, TS performed RNA extraction. SK and PW established and performed the IC-MS analysis. BL, TS, JMH, and ME analyzed transcriptomic data. LW performed and analyzed microfluidic experiments. JMH, TS, LW, and ME drafted the manuscript, with editing by SK, DK, and APMW. DK, APMW, and ME supervised the project and acquired funding. All authors read and approved the final version of the manuscript.

## SUPPLEMENTARY MATERIAL

The following materials are available:

**Supplementary Figure S1:** Close-up on the 1-hour and 2-hour night phases as displayed in Figure 2C.

**Supplementary Figure S2:** Global transcriptional changes in Δ*glgC* mutants at EoN versus EoD.

**Supplementary Figure S3.** Daytime-specific transcript accumulation of photosynthetic light reaction genes.

**Supplementary Figure S4.** Transcriptional responses of Δ*glgC* mutants compared to WT grown in constant light.

**Supplementary Figure S5.** Heatmap for metabolic changes between EoN and EoD samples.

**Supplementary Figure S6.** Relative expression levels of genes involved central carbon metabolism.

**Supplementary Figure S7.** Distribution of secondary site mutations across *Synechocystis* WT and Δ*glgC* mutants.

**Supplementary Figure S8.** Differential gene expression of stress and photoprotection-related genes.

**Supplementary Figure S9.** Relative abundances of nucleotide-sugars detected via IC-MS.

**Supplementary Figure S10.** EoN transcriptional response of genes involved in respiratory electron transport.

**Supplementary Figure S11.** Insertional inactivation of *glgC*.

**Supplementary Table 1:** GO-terms used to select transcripts involved in DNA replication and cell division.

**Supplementary Table 2:** Tools used for the generation of genome assemblies and gff files.

### Supplementary Data File

**Supplementary Dataset S1:** Diurnal RNA-seq data including functional annotation data

**Supplementary Dataset S2:** Constant light RNA-seq data including functional annotation data

**Supplementary Dataset S3:** OD-normalised IC-MS peak data

**Supplementary Dataset S4:** Quantification of adenylates from IC-MS data

**Supplementary Dataset S5:** Identified SSMs in ONT-sequencing data

**Supplementary Dataset S6:** Genes affected by unique SSMs

### Supplementary Videos

**Supplementary Video 1:** Video of a growing *Synechocystis* Δ*glgC*-1 cell in a monolayer growth chamber under constant illumination.

**Supplementary Video 2:** Video of a non-growing *Synechocystis* Δ*glgC*-1 cell in a monolayer growth chamber under constant illumination.

**Supplementary Video 3:** Growth of *Synechocystis* WT in a monolayer growth chamber under diurnal conditions.

**Supplementary Video 4:** Growth of *Synechocystis* Δ*glgC*-1 in a monolayer growth chamber under diurnal conditions.

**Supplementary Video 5:** Growth of *Synechocystis* Δ*glgC*-6 in a monolayer growth chamber under diurnal conditions.

## Notes

### Competing Interest Statement

The authors have declared no competing interest.

