## Supplementary Material for "Glycogen deficiency impairs diurnal energy metabolism and cell division in *Synechocystis*"

*Authors contributed equally

Corresponding author:

**SUPPLEMENTARY FIGURES**

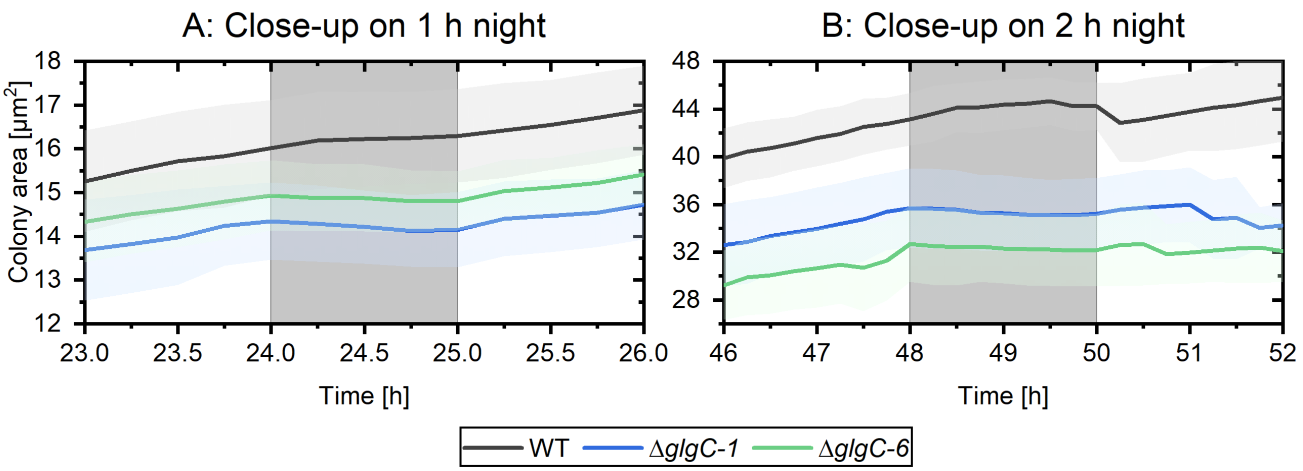


**Supplementary Figure S1.** Close-up on the 1-hour and 2-hour night phases as displayed in Figure 2C. A) During the 1-hour night phase the colony area differences (before – after) were 16.02 – 16.30 $\mu m^2$ (WT), 14.94 – 14.81 $\mu m^2$ (Δ*glgC*-1), and 14.35 – 14.15 $\mu m^2$ (Δ*glgC*-6). B) During the 2-hour night phase the colony area differences (Before – After) were 43.15 – 44.25 µm (WT), 35.71 – 35.21 $\mu m^2$ (Δ*glgC*-1), and 32.70 – 32.19 $\mu m^2$ (Δ*glgC*-6).

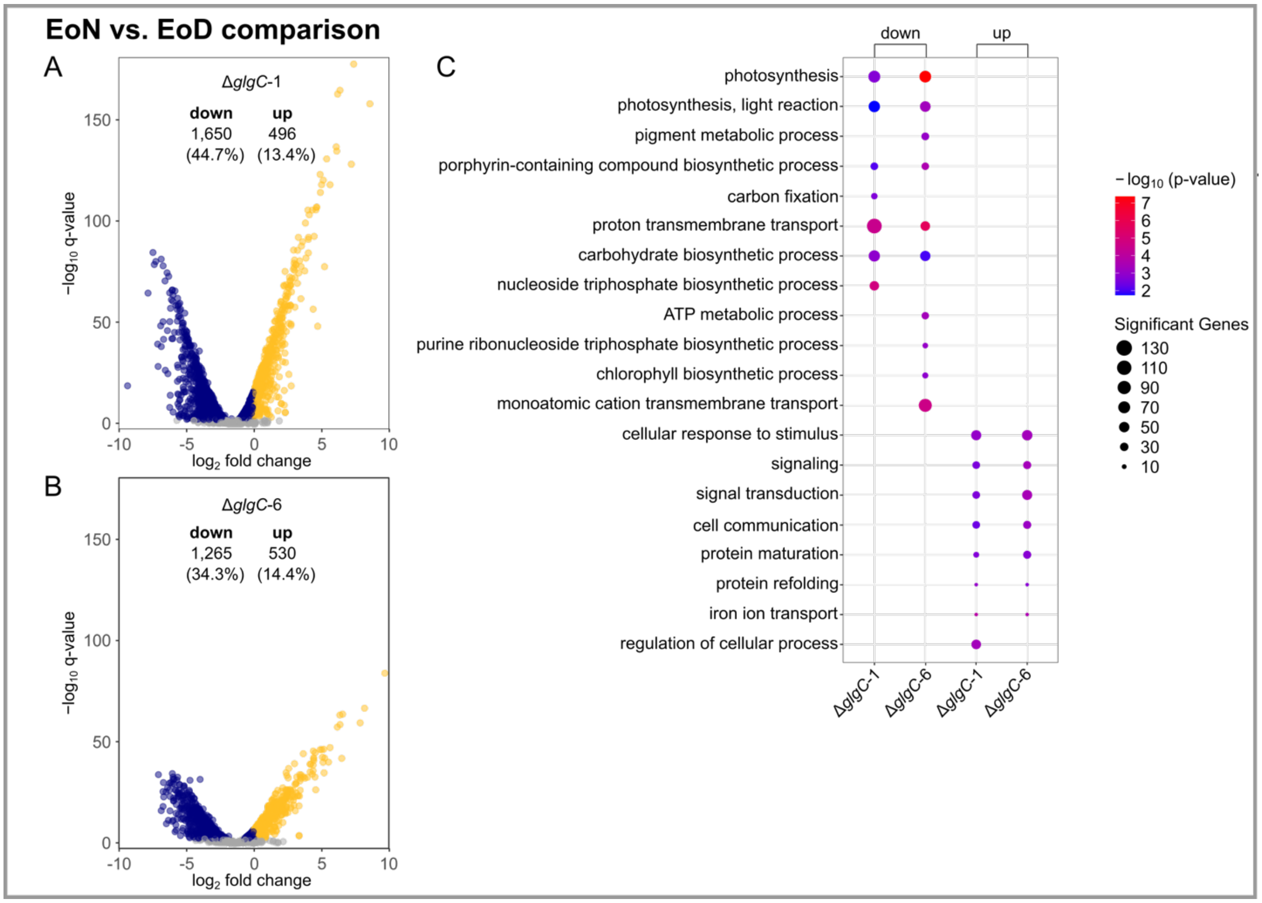


**Supplementary Figure S2.** Global transcriptional changes in Δ*glgC* mutants at EoN versus EoD. (A) Volcano plot of differentially expressed genes (DEGs, $q \leq 0.01$) in Δ*glgC*-1 comparing EoN to EoD. Numbers indicate up- and downregulated genes and their percentage of total coding genes (3,691). (B) Volcano plot of differentially expressed genes (DEGs, $q \leq 0.01$) in Δ*glgC*-6 comparing EoN to EoD. (C) GO term enrichment of DEGs ($q \leq 0.01$) in Δ*glgC* mutant cells. Top 20 biological process GO terms with $p \leq 0.05$ are shown.

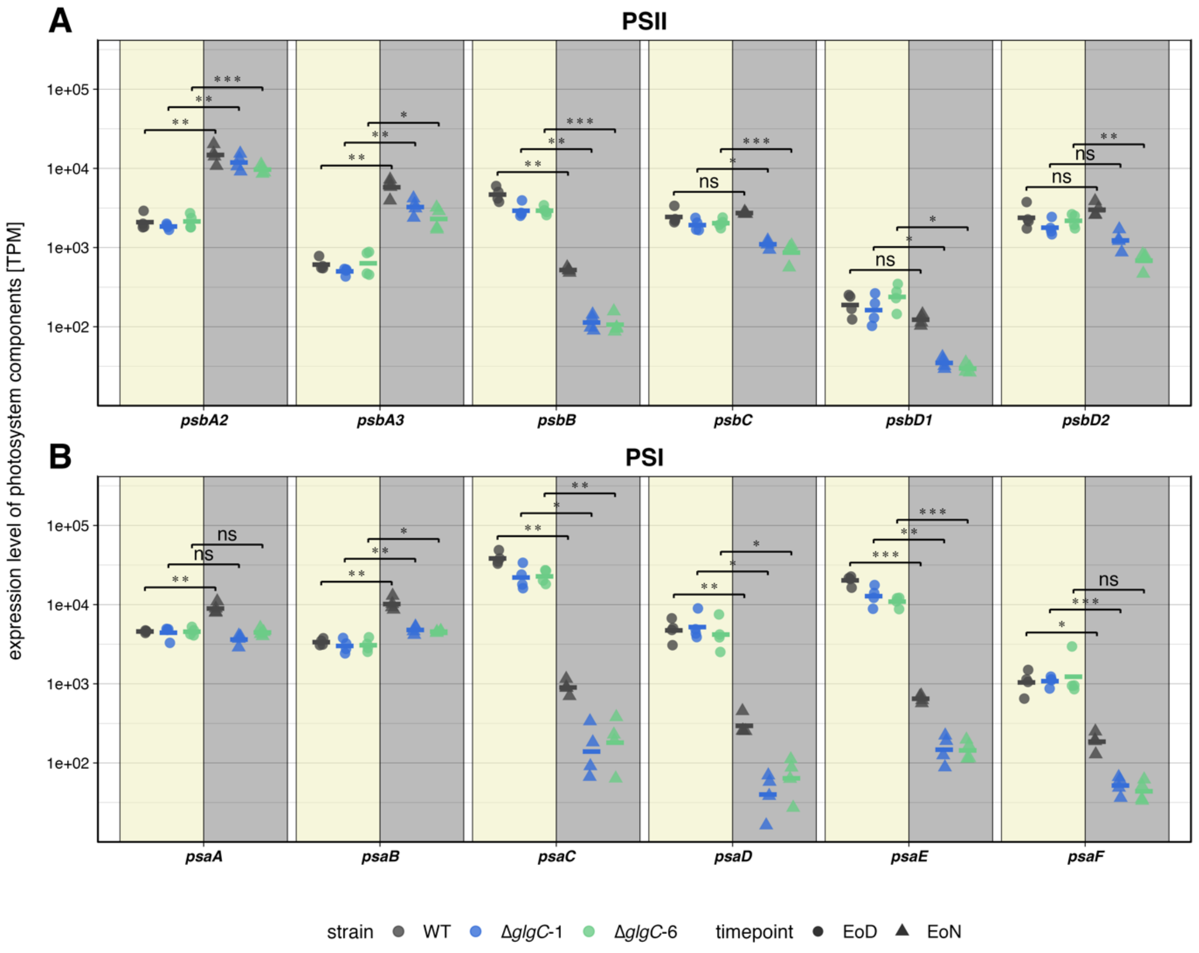


**Supplementary Figure S3.** Daytime-specific transcript accumulation of photosynthetic light reaction genes. (A) TPM values of PSII-encoding genes. (B) TPM values of PSI-encoding genes. Data represent the average of four biological replicates. Individual TPM values are provided in Supplementary Data File 1. Asterisks denote statistical significance between EoN and EoD as calculated with the Student's t-test (*: $p \leq 0.05$, **: $p \leq 0.01$, ***: $p \leq 0.001$).

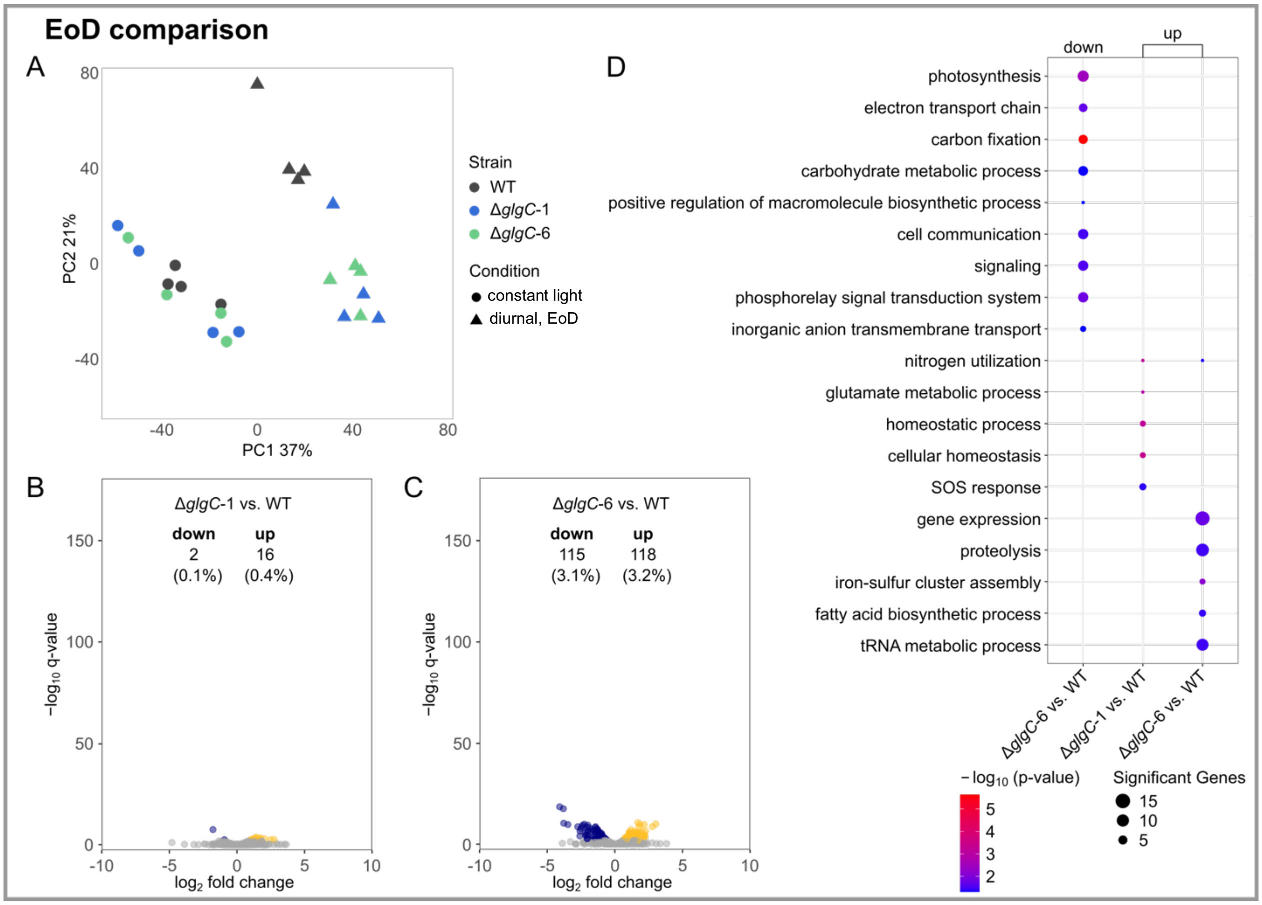


**Supplementary Figure S4*.*** Transcriptional responses of Δ*glgC* mutants compared to WT grown in constant light. (A) PCA of transcript abundances in WT, Δ*glgC*-1, and Δ*glgC*-6 grown under constant light or diurnal regimes, sampled at the end of day (EoD). Transcript per million (TPM) values were used for the analysis. (B) Volcano plot of DEGs ($q \leq 0.01$) in Δ*glgC*-1 versus WT grown under constant light. For all Volcano plots, numbers of significantly up- and downregulated genes and their percentage of total coding genes (3,691) are given. (C) Volcano plot of DEGs ($q \leq 0.01$) in Δ*glgC*-6 versus WT at EoD. (D) GO term enrichment of DEGs ($q \leq 0.01$) for both Δ*glgC* lines. Shown are enriched GO terms (biological processes) with a $p$-value ≤ 0.05.

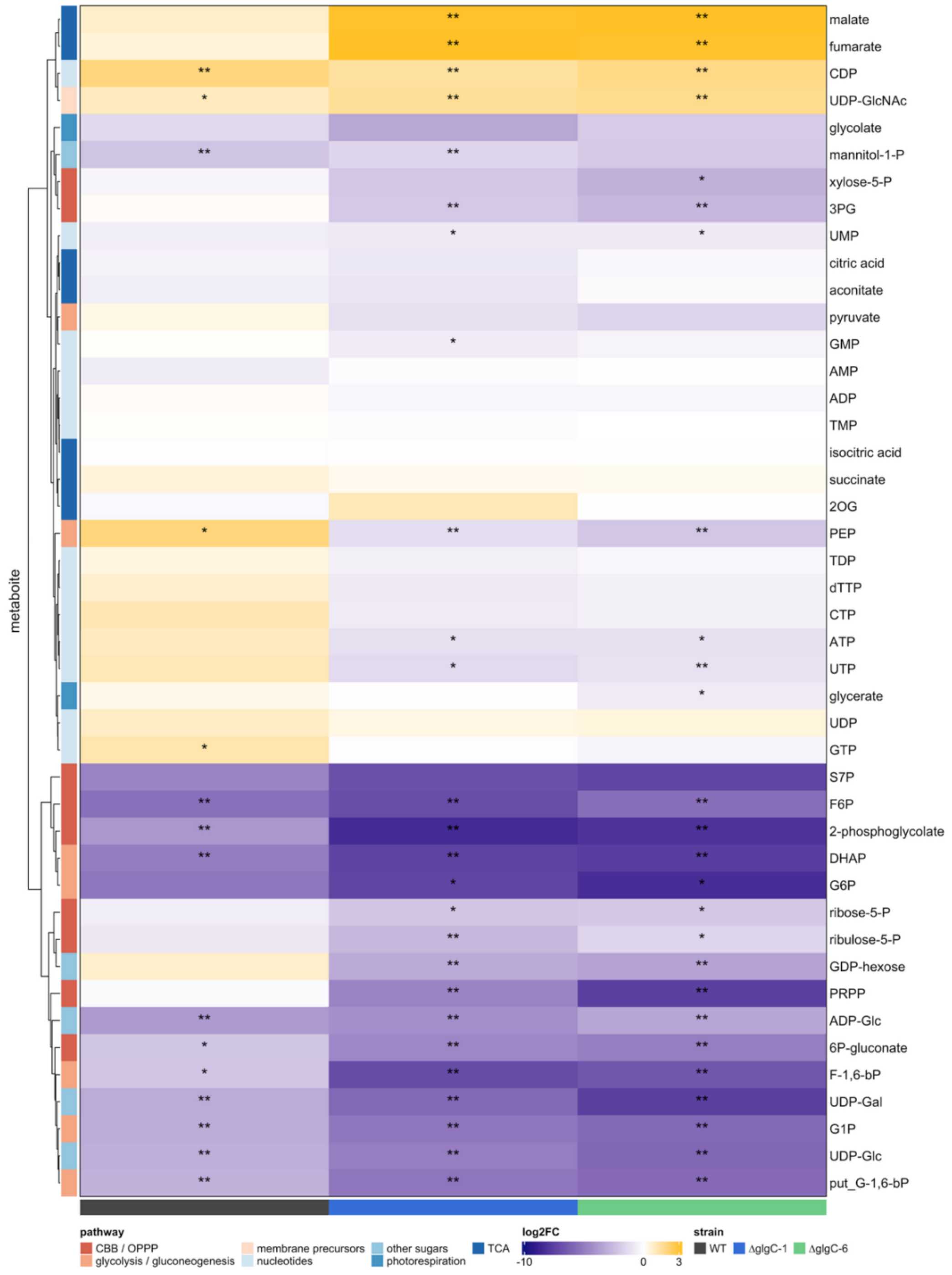


**Supplementary Figure S5:** Heatmap for metabolic changes between EoN and EoD samples. Mutant and WT metabolite levels were determined using IC-MS, median $\log_2$-fold-changes of normalized peak areas for EoN to EoD are displayed. Stars indicate *p*-values of a Wilcoxon-rank test, with * = $p < 0.05$, ** = $p < 0.01$.

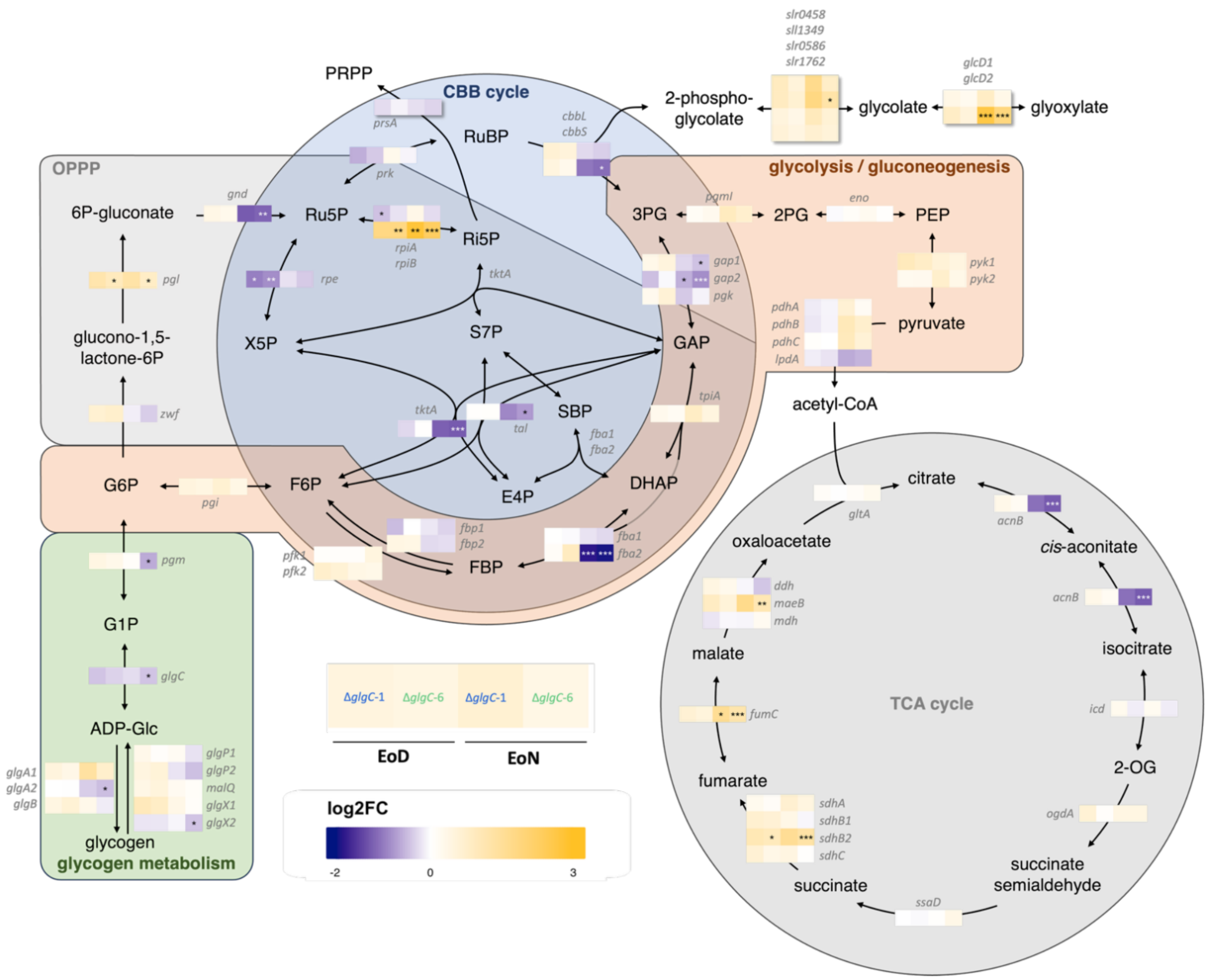


**Supplementary Figure S6:** Relative expression levels of genes involved central carbon metabolism. Log2-fold-changes in expression as determined using edgeR for mutants relative to the WT are shown. Stars indicate *p*-value of a two-sided t-test (* = $p < 0.05$, ** = $p < 0.01$, *** = $p < 0.001$) of n = 4 biological replicates.

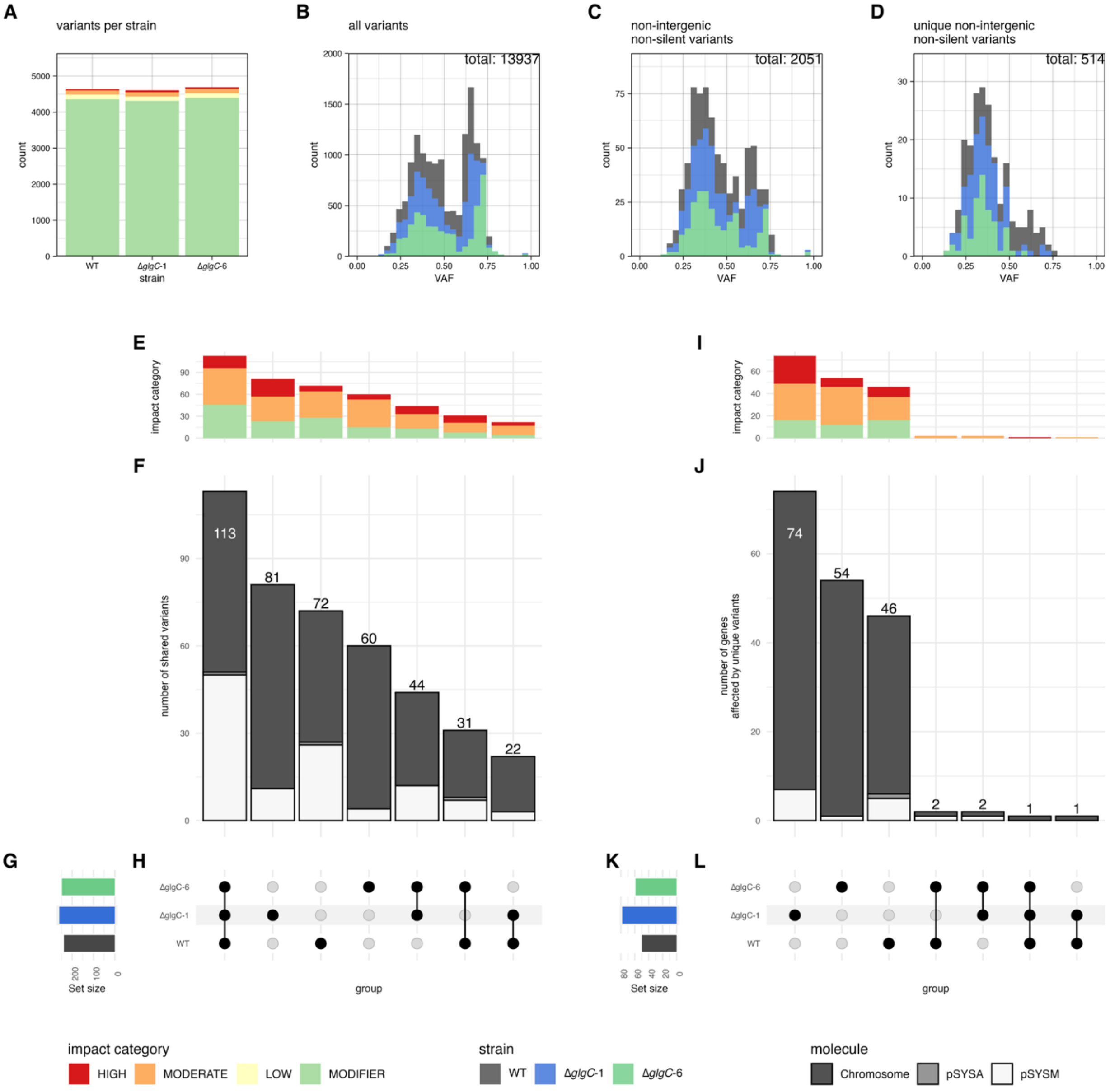


**Supplementary Figure S7:** Distribution of secondary site mutations across *Synechocystis* WT and Δ*glgC* mutants. A) Number of variants found in each strain displaying the impact categories assigned by SnpEff. "HIGH" impact represents disruptions of coding sequences via frame-shifts, early stop codons and gene fusion, "MODERATE" amino-acid changes in protein coding regions, "LOW" synonymous changes at protein level and "MODIFIER" variants outside coding regions. B-D) Histogram of variant allele frequency (VAF) distribution, coloured by strain where the respective variant was found. B) Includes all variants identified; C) non-silent variants in coding sequences (CDS) and mutations < 250 bp upstream of a CDS, considered as promoter regions; D) putative *de-novo* variants occuring only in one strain. E-H) Upset plot displaying the distribution of variants with putative impact (as assigned in C) across genotypes. I-L) Genes affected by *de-novo* variants.

E + I display the impact distribution as assigned by SnpEff, F + J show the number of variants / genes found in each intersect displayed in H + L respectively, with G&L displaying the total number of variants / genes found in each strains.

**Variant calling of mutants**

To examine strain specific differences observed in metabolic and transcriptomic experiments we investigated the occurrence of secondary site mutations in *Synechocystis* WT and Δ*glgC* mutant strains. Long-read sequencing reads obtained from ONT-sequencing were used to assemble genomes and re-aligned to the WT genome (https://www.ncbi.nlm.nih.gov/datasets/genome/GCF_000009725.1/.) using minimap2 (Li, 2021). Deepvariant (Poplin *et al.*, 2018) was used to call variants from the aligned reads. Variant effects were assigned using SnpEff (Cingolani *et al.*, 2012) with custom databases built from the genome annotation. Due to the polyploidic nature no additional variant allele frequency cutoff was set to not lose non-segregated mutations, which could still have biological effects. Analysis and visualization were performed using custom R scripts.

Supplementary Figure S7A shows that most variants were found in intergenic regions, as these are not subject to purifying selection. Supplementary Figure S7B shows the variant allele frequency (VAF) distribution across all called variants. Most variants are present in less than 50% of all reads covering that position, while a notable subset of variants was called in 60-75% of all covering reads. Almost no variants appear to be fully segregated. Variants were then filtered to those with a potential biological impact by selecting variants either ≤ 250 bp upstream of a coding sequence to capture potential changes as a result of altered promoter sequences or causing non-silent variants in coding sequences (Supplementary Figure S7C), which notably does not alter the VAF distribution although 85% of variants were discarded.

Supplementary Figures S7A and S7B-D show that when re-aligning WT reads to the WT genome, the number of identified mutations is comparable to those found in the mutants. As the mutant strains were derived from the WT they are likely to have inherited these mutations. To separate inherited effects from newly acquired variants we assigned unique IDs to variants. Each identical base change at the same position relative to the nearest coding sequence was treated as one variant originating from the same mutation event. Supplementary Figure S7E-H show how variants with a putative effect on gene functionality (in CDSs and non-silent or ≤ 250 bp upstream of a CDS) are distributed across the three genomes. The largest subset of variants can be found in all strains and the rate of mutation does not differ notably between strains, though Δ*glgC*-1 appears to contain more high-impact SSMs than Δ*glgC*-6 and the WT. However, *Synechocystis*' native 100 kbp pSYSM plasmid is overrepresented in the data and appears to have a much higher mutation rate than the chromosome or other plasmids. Unique variants identified in only one strain are much less prevalent (Supplementary Figure S7I-L). To investigate whether these unique mutations indicate adaptive strategies, genes affected by such variants were compared across all strains (Supplementary Figure S7I-L). Here Δ*glgC-1* presents a higher number of genes affected by independent variants than the WT or Δ*glgC-6* do. The overrepresentation of pSYSM is reduced. If the selective pressure was high enough to drive the emergence of secondary site mutations to cope with glycogen-deficiency we would expect that similar genes be affected in the mutants that are not found in the WT, which is neither supported by our findings presented in Supplementary Figure S7F nor S7J. This is not unexpected, as during segregation the mutants were maintained in constant light, where glycogen deficiency has a comparatively low fitness impact. The complete dataset including all called SSMs can be found in Supplementary Dataset S5, and the full list of genes affected by unique variants in Supplementary Dataset S6.

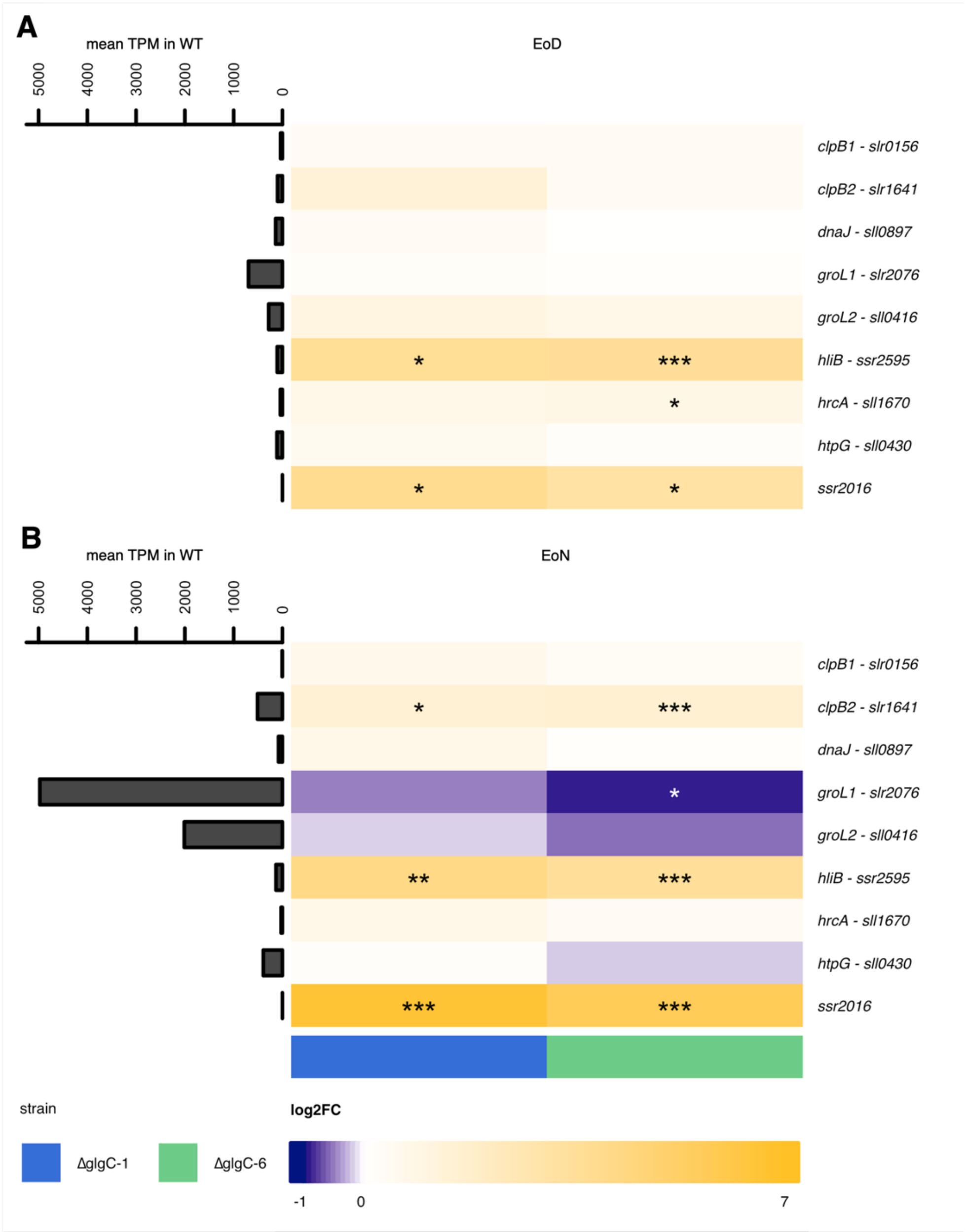


**Supplementary Figure S8:** Differential gene expression of stress and photoprotection-related genes. Expression of selected genes in the mutant lines is compared with expression in the WT at (A) EoD and (B) EoN.

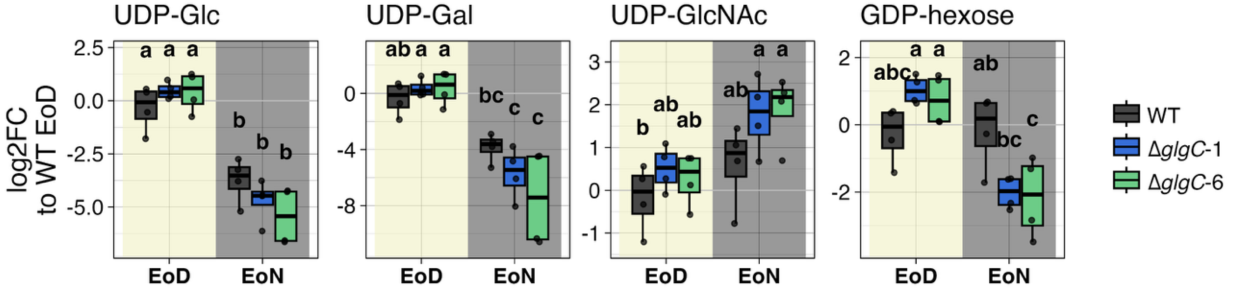


**Supplementary Figure S9:** 1Relative abundances of nucleotide-sugars detected via IC-MS. Log$_2$-fold-changes were calculated from normalised peak areas compared to median normalised peak area in the WT at EoD. Compact letter display indicates significant differences between groups as determined by a Kruskal-Wallis test and Tukey post-hoc analysis of n = 4 biological replicates.

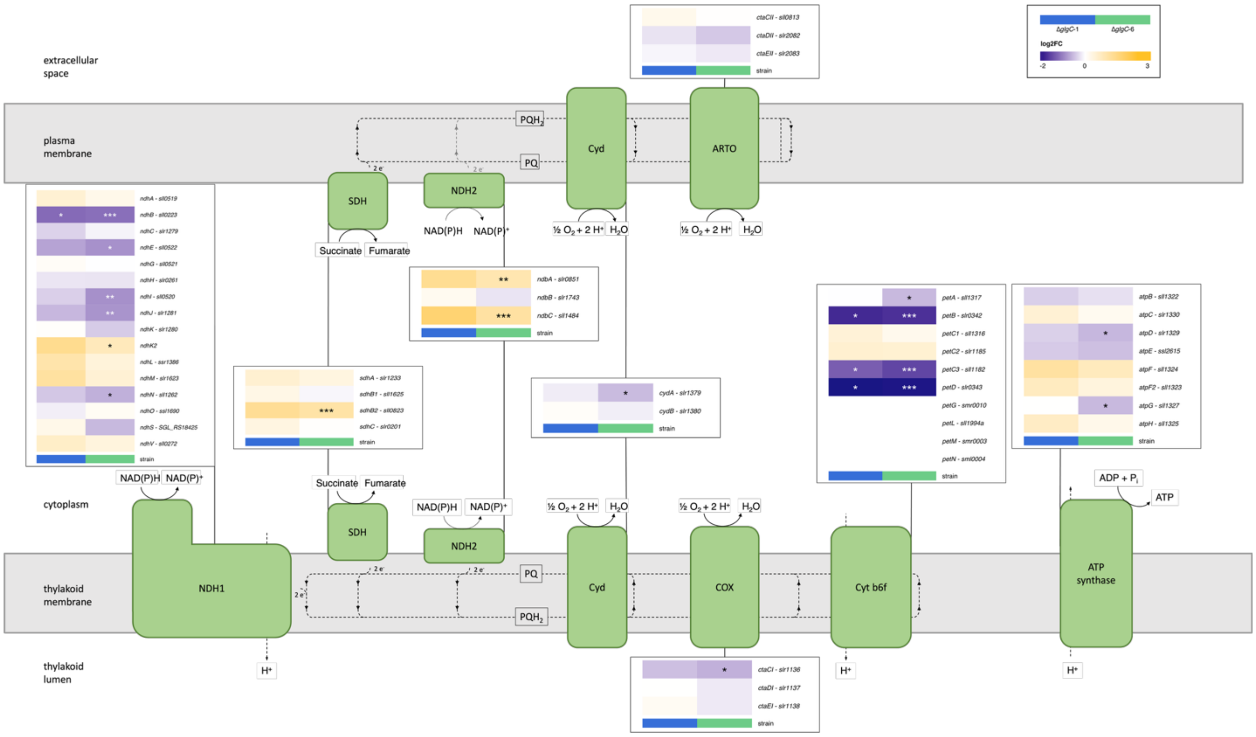


**Supplementary Figure S10:** EoN transcriptional response of genes involved in respiratory electron transport. Function of the corresponding protein complexes is briefly summarized in the figure, adapted from (Lea-Smith et al., 2016). Genes were selected based on association with the corresponding protein complexes in CyanoCyc v29.0.

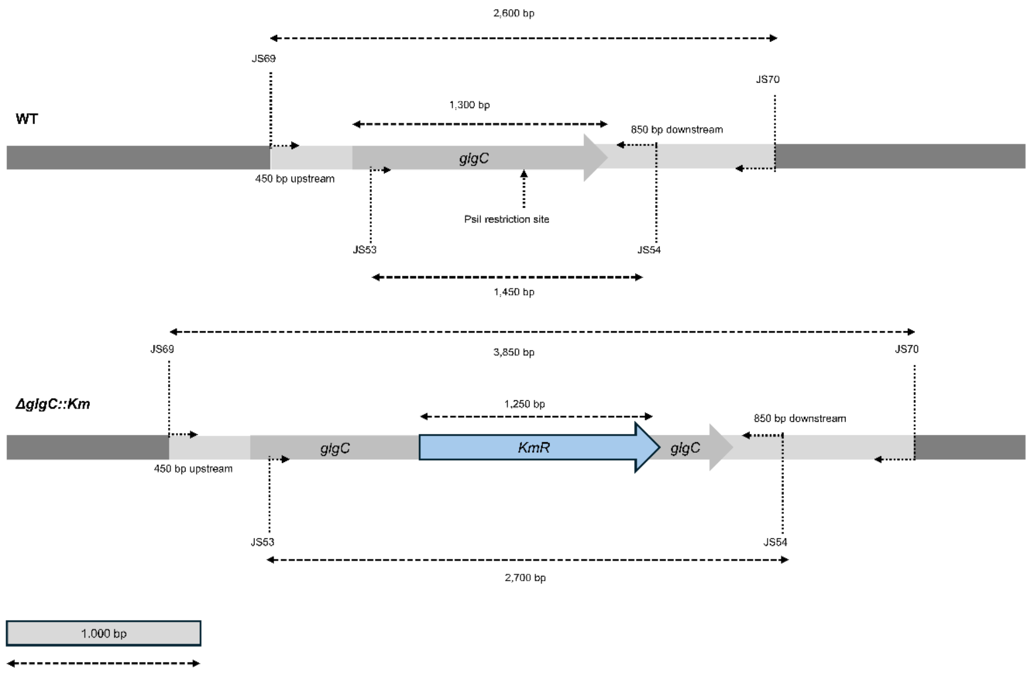


**Supplementary Figure S11:** Insertional inactivation of *glgC* (*slr*1176) to generate Δ*glgC* mutants. A plasmid was constructed by cloning of the *glgC* gene along with up- and downstream regions into a vector. A kanamycin resistance cassette (*KmR*) was inserted into the PsiI site. By homologous recombination, the kanamycin resistance cassette is inserted into the *glgC* gene, thereby insertionally inactivating the open reading frame. Full segregation of the mutants` genome was checked with the primer pair JS53/JS54.

**SUPPLEMENTARY TABLES**

**Supplementary Table 1:** GO-terms used to select transcripts involved in DNA replication and cell division. The CyanoCyc v29.0 was filtered for all genes matching at least one of these GO terms.

| Category, Figure | GO-Term | Description |
|---|---|---|
| DNA replication, Figure 8C | GO:0006259 | DNA metabolic process |
| | GO:0055133 | DNA replication (obsolete) |
| | GO:0006261 | DNA templated DNA replication |
| | GO:0006269 | DNA replication, synthesis of primer |
| | GO:0006271 | DNA strand elongation involved in DNA replication |
| | GO:0045005 | DNA-templated DNA replication maintenance of fidelity |
| | GO:0006270 | DNA replication initiation |
| | GO:0006260 | DNA replication |
| Cell Division, Figure 8D | GO:0007049 | cell cycle |
| | GO:0051301 | cell division |
| | GO:0051302 | regulation of cell division |
| | GO:0051381 | positive regulation of cell division |
| | GO:0051782 | negative regulation of cell division |

**Supplementary Table 2:** Tools used for the generation of genome assemblies and gff files.

| Software | Version | Reference |
|---|---|---|
| samtools | 1.23 | (Danecek *et al.*, 2021) |
| porechop | 0.2.4 | (Wick *et al.*, 2017) |
| seqkit | 2.12.0 | (Shen, Sipos and Zhao, 2024) |
| flye | 2.9.6 | (Kolmogorov *et al.*, 2019) |
| racon | 1.5.0 | (Vaser *et al.*, 2017) |
| medaka | 2.1.1 | (Oxford Nanopore Technology, 2018)5/21/26 5:33:00 PM |
| berokka | 0.2.3 | (Seemann, 2018) |
| minimap2 | 2.30 | (Li, 2018, 2021) |
| circlator | 1.5.5 | (Hunt *et al.*, 2015) |
| quast | 5.3.0 | (Mikheenko *et al.*, 2023) |
| BUSCO | 6.0.0 | (Stanke *et al.*, 2008; Camacho *et al.*, 2009; Hyatt *et al.*, 2010; Eddy, 2011; Mirarab, Nguyen and Warnow, 2012; Levy Karin, Mirdita and Söding, 2020; Li, 2023; Tegenfeldt *et al.*, 2025) |
| Liftoff | 1.6.3 | (Shumate and Salzberg, 2021) |

### SUPPLEMENTARY METHODS

#### Constant Light Experiment

Pre-cultures of WT, Δ*glgC*-1, and Δ*glgC*-6 were grown under standard cultivation conditions. After the $OD_{750}$ was sufficiently high, the cultures were harvested via centrifugation (20 minutes at 3,000 x *g*). The cells were washed once with fresh BG11 medium and resuspended in 15 mL BG11 medium. The main cultures were adjusted to an $OD_{750}$ of 0.7 in 70 mL of BG11. The cultivation was performed in the Multi-Cultivator MC-1000-OD (Photon Systems Instruments, Drásov, Czech Republic) under constant illumination of 100 µmol photons $m^{-2}$ $s^{-1}$ at 30 °C. After 36 h, two 10 mL samples of each culture were collected and centrifuged at 3,000 x *g* for 10 min at 4 °C and the supernatant was discarded. Pellets were stored in the -80 °C freezer. RNA was extracted from one 10 mL sample and sequenced via RNA-seq and then analyzed as described in the material and methods section.

**Long-read sequencing of *Synechocystis* strains**

Cultures of WT, Δ*glgC*-1, and Δ*glgC*-6 were grown in the Multi-Cultivator MC-1000-OD. The cultures were harvested, and 50 mL of culture volumes was centrifuged at 3,000 x *g* for 20 min. The supernatant was discarded and the gDNA extraction was performed as described in (Theune *et al.*, 2026). The library preparation was performed with the LSK114 Ligation Sequencing KIT. The sequencing run was performed on the PromethION (MinKNOW 23.11.7, Bream 7.8.2, Configuration 5.8.6, MinKNOW Core 5.8.6) and a minimum 100 x coverage was achieved. Bases were called using Dorado 7.2.13.

***De-novo* assembly of *Synechocystis* genomes**

All tools used for the assembly of the genomes are listed in Supplemental Table 2. Adapters were trimmed with porechop (Wick *et al.*, 2017). Genome was assembled with flye (Kolmogorov *et al.*, 2019) and polished. Additional polishing was performed with racon (Vaser *et al.*, 2017) and medaka (Oxford Nanopore Technology, 2018). Overlaps in the sequences were trimmed with berroka (Seemann, 2018). Sequences were linearized with circulator (Hunt *et al.*, 2015). Assembly qualities were checked with quast (Mikheenko *et al.*, 2023) and BUSCO (Tegenfeldt *et al.*, 2025). Genome annotation was generated by using liftoff (Shumate and Salzberg, 2021) to map the refseq gff file of the genome assembly ASM972v1 (https://www.ncbi.nlm.nih.gov/datasets/genome/GCF_000009725.1/.) to our WT assembly. The raw reads along with the assembled genome with their corresponding annotated features were uploaded to ENA and are available under the BioProject accession number PRJEB112054.
